# Re-emergence of orientation coding in primate IT cortex and deep networks reveals functional hubs for visual processing

**DOI:** 10.1101/2025.10.23.684240

**Authors:** Behnam Karami, Tarana Nigam, Sandrin S. Plewe, Caspar M. Schwiedrzik

## Abstract

The primate visual system is a hierarchical network of brain areas that transform retinal inputs into rich percepts. According to efficient neural coding principles, basic visual features such as orientation, which are already represented in early visual areas, should not be redundantly encoded in higher areas like V4 and inferotemporal (IT) cortex. We tested this hypothesis by quantifying orientation representation throughout the entire visual stream using functional Magnetic Resonance Imaging (fMRI) and multivoxel pattern analysis (MVPA) in macaque monkeys watching full-field gratings. Contrary to predictions from efficient coding, we found robust orientation information along the entire visual ventral stream from V1 to IT cortex. Orientation information was distributed in cortical and representational space, especially in IT. Voxels with multivariate responsiveness to faces, objects, and bodies carried more orientation information than highly category-selective voxels. They also displayed stronger connectivity within IT and with upstream areas V1–V4. These findings suggest the existence of functional “hubs” that integrate low- and high-level features through enhanced connectivity. This phenomenon generalized to deep convolutional neural network models with high brain similarity, underscoring the importance of hub-like units in hierarchical processing of complex inputs. Together, our results show that orientation re-emerges in IT cortex not through redundancy but as a computational byproduct of functional diversity and connectivity, propounding the existence of “hub” neurons that may support flexible, integrative visual processing beyond object recognition.

## Introduction

The primate visual system is a complex network of brain areas that operates on retinal inputs to generate rich percepts across multiple stages of processing. In cortex, this computation is thought to begin with the extraction and representation of low-level visual features, such as orientation, color, luminance, contrast, etc., in V1. Processing then proceeds to mid-level features like curvature and depth in areas V3 and V4, and finally culminates in high-level category representations, such as faces, bodies, and objects in inferotemporal (IT) cortex (Felleman & Van Essen, 1991; Kravitz et al., 2013). This hierarchical framework has garnered significant support in systems neuroscience, particularly with the advent of Deep Convolutional Neural Networks (DCNNs) (Barlow, 1961; Burnston & Haueis, 2021).

Efficient neural coding suggests that low-level features should not be redundantly represented in higher-level visual areas of the visual processing hierarchy (Barlow, 1961; Burnston & Haueis, 2021). Instead, such features are thought to be dropped from higher stages of processing as representations become increasingly complex and invariant (Riesenhuber & Poggio, 1999; Ullman, 2007). However, other theories suggest that information from lower levels is not discarded but instead preserved across sequential transformations in the visual processing hierarchy, allowing for data-efficient reuse for future inference (Higgins et al., 2022). Indeed, empirical evidence suggests that certain low-level features—such as color, luminance, and contrast—are represented in high-level visual regions like V4 and IT, alongside their encoding in lower-level areas (Ai et al., 2025; Lafer-Sousa & Conway, 2013; M.-L. Liu et al., 2024; Y. Liu et al., 2020; Marquardt et al., 2018; Namima et al., 2014; Taylor & Xu, 2022; Vladusich et al., 2006). This redundancy may be important to enable, e.g., rapid and generalizable plasticity when dealing with highly variable environments (Ahissar & Hochstein, 2004; Manenti et al., 2023). Whether redundancy reduction or a distributed form of coding for low level features, respectively, constitute a fundamental principle of visual processing remains an open and highly debated question (Bracci et al., 2017; Reddy & Kanwisher, 2006). In this study, we investigate whether distributed encoding extends to the feature of orientation, which has not yet been systematically explored in this context.

Orientation is a fundamental visual attribute and a key building block for the reconstruction of object shape. Beginning with Hubel and Wiesel’s pioneering work on the cat’s visual cortex, where they first identified neurons responsive to oriented bars (Hubel & Wiesel, 1962), a lot has been revealed about the organization and structure of orientation representations in early visual areas, most prominently pin-wheel columns in V1 and thick-and-thin stripes in V2 (Dow, 2002; Felleman et al., 2015). However, much less is known about how orientation is represented in higher-level visual areas, particularly beyond V4 (Desimone et al., 1985; Vanduffel et al., 2002). This may be due to the challenges of recording from the entire visual cortex, difficulties in eliciting responses to simple stimuli such as gratings in early recordings of IT cortex (Gross, 2008), and/or the strong influence of the traditional hierarchical processing model.

We sought to systematically investigate orientation representations across the entire visual ventral stream using functional magnetic resonance imaging (fMRI) in awake macaque monkeys to draw a direct link with well-established electrophysiological and neuroimaging findings on orientation processing available in this species. Given the coverage and spatial resolution that fMRI affords, this technique is ideally suited for capturing local as much as distributed representations in the brain. We find that orientation representations extend over the entire visual ventral stream, but in a distinct, distributed form that differs from other visual features. In particular, the spatial arrangement of orientation representations across the hierarchy exhibits an hourglass-shaped organization, where small clusters in early visual areas (V1-V2) converge into larger clusters in mid-level areas (V3-V4), before diverging again into numerous clusters in IT. Orientation representations, especially in areas beyond V4, thus fundamentally differ from the highly localized, patchy representation of color and visual object categories (Bao et al., 2020; Lafer-Sousa & Conway, 2013; Tsao et al., 2006).

To further explore the potential role of orientation representations in high-level areas, we examined their relationship with a well-established function of IT cortex—visual object category representation. Interestingly, we find that voxels showing low category selectivity for faces, objects, and bodies carry more orientation information than those responding preferentially to a single category. This finding replicates not only in the brain, but also across multiple DCNN architectures. Furthermore, the orientation-responsive, non-category selective voxels display enhanced functional connectivity with other voxels within IT and with upstream visual areas compared to strongly category selective voxels, suggesting a privileged role in distributing information in a category-agnostic way.

In light of these results, we propose that orientation representations in IT cortex may arise from enhanced connectivity with early visual areas, the primary source of orientation information in cortex. Furthermore, we suggest the existence of “hubs” in IT cortex — clusters that receive upstream input with privilege, process visual information non-selectively, and distribute this information to more specialized neural clusters.

## Results

### Presence of orientation information along the ventral stream

Orientation is represented in a mesoscale, columnar manner across early visual cortex below the resolution of fMRI at 3T. However, multivoxel pattern analysis (MVPA) enables decoding of orientation information by analyzing spatial patterns across multiple voxels (Harrison & Tong, 2009; Haynes, 2015; Kamitani & Tong, 2005). Leveraging this technique, we computed orientation information across the entire visual ventral stream and several control regions in the macaque brain. To that end, we acquired blood oxygenation level-dependent (BOLD) signals at 3T in two male adult monkeys (monkey R, and P) while they fixated on full-field Gabor gratings showing one of two orthogonal orientations at a time (45° or 135°; Fig. 1A). We decoded orientation from each brain area using Support Vector Machines (SVM) trained on BOLD activity patterns of the entire region (Fig. 1B).

**Fig. 1.**
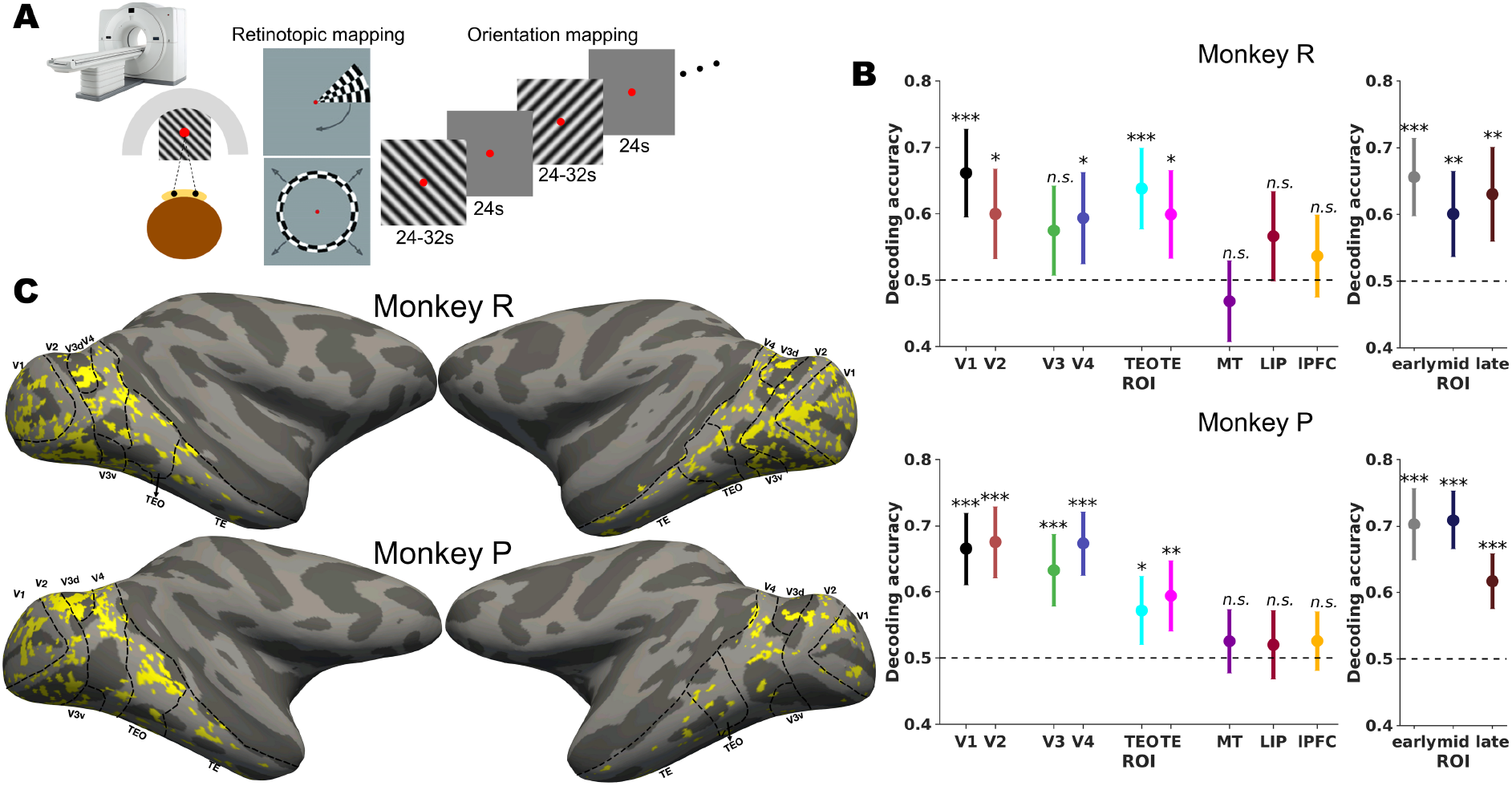
Experimental paradigm and multivoxel pattern analysis (MVPA) results. **A.** fMRI data were acquired from two macaque monkeys during passive fixation. Each monkey underwent retinotopic mapping (polar angle and eccentricity) to define visual regions of interest (ROIs), followed by orientation mapping using two diagonal grating orientations to assess orientation sensitivity across the entire cortex. **B.** Orientation information quantified via MVPA across ROIs. A linear support vector machine (SVM) classifier was trained to distinguish the two diagonal orientations based on multivoxel patterns. Error bars represent standard deviation across 100 cross-validation splits. Decoding accuracy was tested against a null distribution generated by shuffling orientation labels, using a one-sided permutation test (10,000 permutations, FDR-corrected across ROIs; statistics in Results). Asterisks: *** p < 0.001, ** p < 0.01, * p < 0.05; n.s., not significant. **C.** Orientation-responsive voxels defined based on contrast t-statistics (*p* < 0.05) and projected onto inflated surface maps for both monkeys as indicated in yellow. Visual ROI boundaries are outlined with dashed lines. The image of the MRI in A is reproduced with permission from Adobe.

Decoding accuracy was significantly above chance in all areas along the ventral stream in both animals (with the exception of V3): in monkey R, mean decoding accuracy (100 cross- validation folds, one-sided permutation tests, FDR-corrected) was 66.1% (± SD 6.6%, p < 0.001) in V1, 60.0% (± SD 6.8%, p = 0.0180) in V2, 57.5% (± SD 6.7%, p = 0.0604) in V3, 59.4% (± SD 6.9%, p = 0.0288) in V4, 63.8% (± SD 6.1%, p < 0.001) in TEO, and 59.9% (± SD 6.6%, p = 0.0180) in TE. In monkey P, mean decoding accuracy was 66.5% (± SD 5.4%, p < 0.0001) in V1, 67.5% (± SD 5.4%, p < 0.0001) in V2, 63.2% (± SD 5.4%, p < 0.001) in V3, 67.3% (± SD 4.8%, p < 0.001) in V4, 57.2% (± SD 5.1%, p = 0.0135) in TEO, and 59.4% (± SD 5.3%, p = 0.0018) in TE. Pooled by processing stage, decoding was significant in early, mid, and late stages in both monkeys: in monkey R, mean decoding accuracy was 65.6% (± SD 5.8%, p < 0.001) in the early, 60.0% (± SD 6.4%, p = 0.0075) in the mid, and 63.0% (± SD 7.1%, p = 0.0023) in the late stage. In monkey P, mean decoding accuracy was 70.3% (± SD 5.4%) in the early, 70.9% (± SD 4.4%) in the mid, and 61.7% (± SD 4.1%) in the late stage (all p < 0.001). These results replicate the well-known involvement of early visual cortex in orientation processing and extend this function well into the anterior temporal lobe.

To verify the specificity and reliability of these effects, we measured orientation decoding in higher-order and dorsal stream areas (MT, LIP, and lateral prefrontal cortex (lPFC)) using the same procedure. Decoding accuracy was not significantly above chance in any of these areas in any of the animals: in monkey R, mean decoding accuracy was 46.8% (± SD 6.1%, p = 0.8010) in MT, 56.6% (± SD 6.7%, p = 0.6040) in LIP, and 53.6% (± SD 6.2%, p = 0.2036) in lPFC. In monkey P, values were 52.5% (± SD 4.8%, p = 0.2173) in MT, 52.0% (± SD 5.2%, p = 0.2430) in LIP, and 52.6% (± SD 4.4%, p = 0.2273) in lPFC. This supports the specificity of the decoding results in ventral visual cortex.

### Representational spaces for orientation along the ventral stream

As we found that orientation information is present throughout the entire visual ventral stream, from V1 to IT (see Fig. 1C for cortical surface projections), we next sought to investigate whether regions differed in their representational format. We hypothesize that V1, as the earliest cortical stage involved in orientation processing and containing numerous highly orientation-tuned neurons, would encode this feature within a select subset of highly selective voxels. These voxels are commonly defined by their strong univariate orientation contrast between the two diagonal orientations, reflecting a more selective and precise representational format (Gotsopoulos et al., 2018; Kriegeskorte et al., 2006; Mur et al., 2009). Of note, such univariate effects on the single voxel level can even be observed in human fMRI at much coarser resolution than the one we employed here (Albers et al., 2018). In contrast, higher-order areas in later stages may not rely on a specialized subset of highly tuned neurons. Instead, they may encode orientation across a broader population of less selective neurons. In this case, orientation information would be more distributed, rather than concentrated, across both highly and less selective voxels, consistent with a global representation format.

To investigate the distribution of orientation information in representational space within and across areas, we employed a decoding approach relating the number of voxels required for accurate decoding to single voxel orientation informativeness (schematic in Fig. 2A bottom): in each area, we ranked voxels by their (univariate) absolute contrast t-statistics (see Methods for details). We trained the SVM classifier starting with the 10 most informative voxels and incrementally added voxels in descending order of univariate information. This produced, for each area, a decoding accuracy curve as a function of voxel proportion, each reaching a maximum at a specific proportion. We defined peakProp as the voxel proportion at which the curve reached its maximum, which serves as our multivariate metric. The corresponding univariate metric, sigProp, was the proportion of voxels in an area reaching significance in the univariate orientation contrast.

**Fig. 2.**
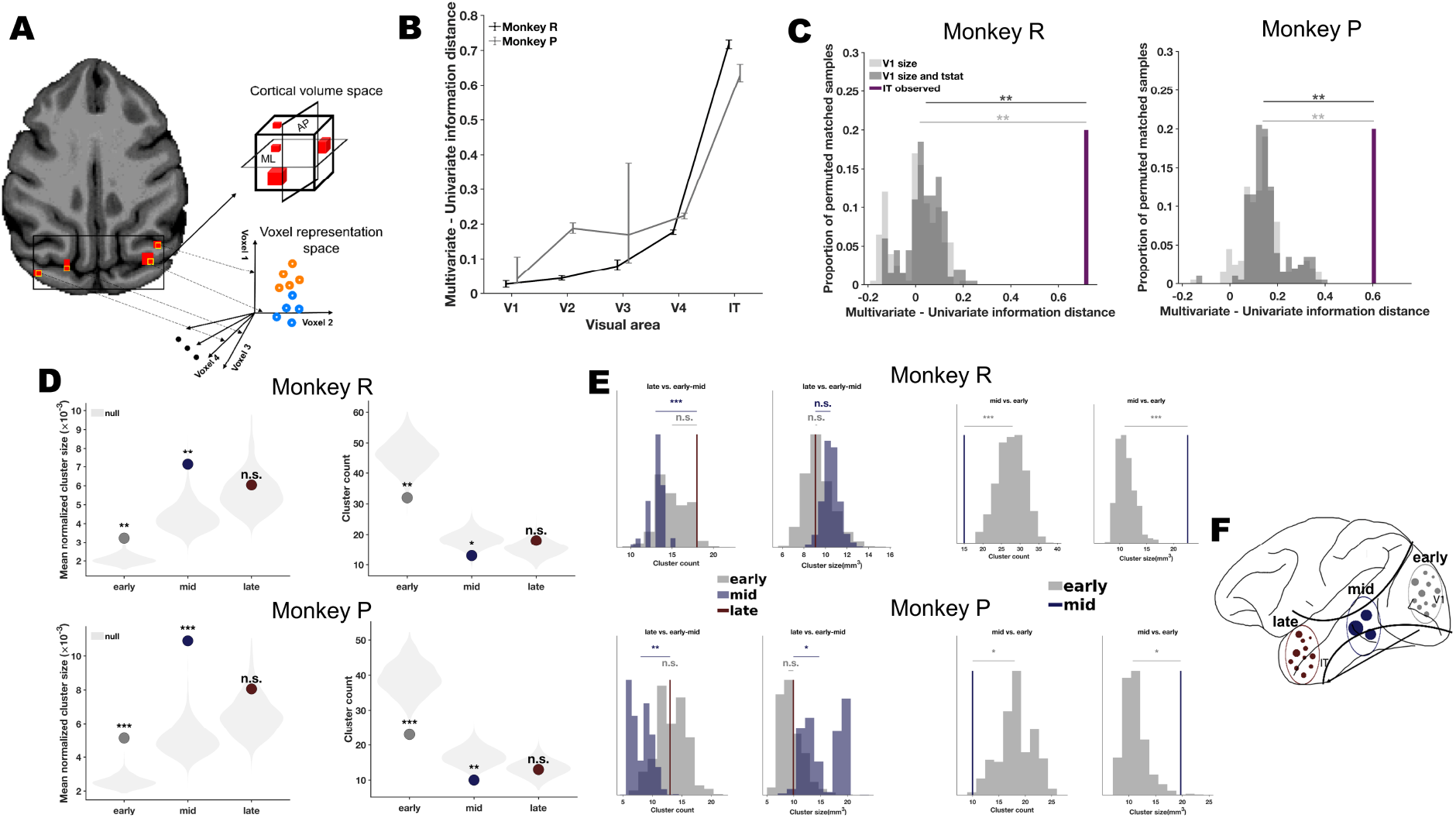
Orientation organization along the visual hierarchy. **A.** Orientation organization was appraised in two domains: anatomical (cortical volume) and representational (multivoxel pattern) space. In cortical space, voxels with high univariate orientation selectivity (*p < 0.05*, based on contrast t-statistics) were clustered in 3D by spatial adjacency. In representational space, voxels were incrementally added in descending order of absolute t-statistic, regardless of anatomical location. **B.** Separation between the multivoxel and univariate descriptions of each area quantified as peakProp - sigProp. peakProp is the proportion of voxels at which the incremental decoding curve reaches its maximum. sigProp is the proportion of voxels (i.e. number voxel included with criteria divided by the total voxels identified in the region) reaching p < 0.05 in the univariate orientation contrast. A near-zero distance means decoding stops improving once the individually selective voxels are in; a large distance means voxels that failed to reach univariate significance continue to add multivariate information. Error bars are 95% intervals from 1000 bootstrap resamples of the classifier repetitions, with the peak re-estimated on each resample. **C.** The observed IT distance (purple) against two null distributions of V1 resamples (200 samples each). *V1 size* subsamples V1 at random to IT’s voxel count. *V1 size and t-stat* additionally matches IT’s full absolute t-stat distribution, so sigProp distance cannot arise from a difference in univariate selectivity or BOLD Signal-to-Noise Ratio (SNR). **D.** Mean normalized cluster size and cluster count per processing stage (markers gray = early, blue = mid, red = late) against the orientation-label permutation null (shaded light gray, 1000 permutations). In every permutation the two orientation labels were swapped at random per block pair, the t-stat map recomputed and used for clustering. **E**. Cluster count and mean cluster size for the late stage against patches drawn from early (gray) and mid (blue), and for the mid stage against patches drawn from early. Patches were matched to the target stage on both volume and number of selected voxels (10,000 samples per source). **F.** Schematic depiction of an hourglass-shaped organization: orientation representations are spatially integrated in cortical space from early to mid stages, then redistributed more diffusely in the late stage. The main redistribution effect was carried more by cluster number and less of cluster size. Statistical comparisons are permutation tests. Asterisks denote *** p<0.001, ** p<0.01, * p<0.05; n.s. denotes not significant.

We then measured the distance between these two metrics and plotted it across visual areas from V1 to IT (Fig. 2B). If all orientation information were contained in a select subset of highly selective voxels, the decoding curve would peak at sigProp, and the remaining voxels would not contribute relevant information. A greater distance between sigProp and peakProp instead indicates a more distributed orientation information profile within the area. We observed a general trend of increasing distance from V1 to IT, most pronounced between V1 and IT (Fig. 2B). This suggests that orientation representations became increasingly dispersed as a function of the visual ventral stream hierarchy.

However, two factors limit a direct comparison between the start- and endpoint of this hierarchy, V1 and IT. First, the two areas differ substantially in size (V1 was more than double the size of IT in both monkeys). Because a given proportion of a larger area corresponds to a larger number of voxels, the V1 curve can reach its maximum at a lower proportion simply because that proportion already affords a high dimensional pattern. Second, given the placement of our receiver coils, contrast t-statistics are possibly inflated in V1 relative to IT, so that fewer IT voxels pass the threshold contributing to sigProp. We controlled for both factors by constructing null distributions from V1 subsamples: one matched to IT in size only (V1size), and one matched to IT in size and additionally to its distribution of absolute t-statistics (V1size and tstat), which equates sigProp by construction. We then compared the observed IT distance against each null using a permutation test (Fig. 2C). The observed distance exceeded both null distributions (monkey R: observed distance = 0.7167; V1size null mean = 0.0056, p = 0.005; V1size and tstat null mean = 0.039 p= 0.005; monkey P: observed = 0.6044; nulls = 0.1402 and 0.1554, p = 0.005 and 0.005), indicating that orientation information is indeed more distributed in IT than in V1, and that this difference is not attributable to area size or to differences in univariate selectivity.

### Spatial organization of orientation information along the ventral stream

Along the visual hierarchy, neurons become increasingly specialized in representing complex objects and exhibit greater spatially clustering in cortical space. For example, millimeter-scale color-selective globs in V4 arise from the finer-scale blob/interblob organization in V1 and V2 (Conway et al., 2007; Roe & Ts’o, 1999; Tanigawa et al., 2010); and domain-specific patches selective for faces, bodies, objects, and color emerge in IT cortex, consistent with mesoscale organization of primate visual system (Bao et al., 2020; Chen et al., 2025; Lafer-Sousa & Conway, 2013; Y. Lu et al., 2018; Tsao et al., 2006; Wang et al., 2024; Zhu et al., 2025). Organizing feature-selective neurons into specialized clusters is thought to facilitate local computations (Douglas & Martin, 2004), improve downstream readout (DiCarlo et al., 2012), enhance the reliability and robustness of neural code (Op de Beeck et al., 2008), and provide focal targets for experience-dependent plasticity and top-down modulations within these specialized clusters (Bichot et al., 2015; Sigala & Logothetis, 2002). Does this form of organization also apply to orientation?

Visual inspection of differential orientation responses projected onto the cortical surface suggests heterogeneous clustering across the visual cortex, with larger clusters in certain regions, such as area V4 (Fig. 1C). To quantitatively examine this observation, we applied a clustering algorithm in volume space, grouping adjacent orientation-selective voxels into clusters (see Methods for details on the algorithm). We quantified the number and mean normalized size of the clusters (i.e., mean cluster size divided by the region size) across three processing stages: early (V1-V2), mid (V3-V4), and late (TEO-TE/IT).

We held the number of voxels entering the clustering procedure fixed across stages. Rather than thresholding each stage independently, we determined the proportion of selected voxels in the late stage and applied the same proportion to the early and mid stages (see Methods). This way, differences in cluster count and size between stages cannot arise from differences in how many voxels survived statistical selection in each stage.

Cluster count and size varied across stages (Fig. 2D). In monkey R, the early stage contained 32 clusters with a mean normalized size of 0.0032, the mid stage 13 clusters (mean normalized size = 0.0071), and the late stage 18 clusters (0.0060). Monkey P showed the same ordering: 23 clusters (0.0051) in the early stage, 10 (0.0109) in the mid stage, and 13 (0.0081) in the late stage. From early to mid stages, clusters thus became fewer and larger, as expected from convergent integration along the visual hierarchy. From mid to late stages, this trend reversed: clusters became more numerous and smaller, with both count and mean normalized size falling between early and mid stage values.

To determine whether these properties reflect orientation-specific spatial structure, we compared each observed value against a null distribution obtained by permuting the orientation labels and recomputing the t-map, holding the number of selected voxels at its observed value in each permutation (1,000 permutations; Fig. 2D). This null preserves the spatial autocorrelation of the data while removing the orientation-specific structure. Departures from it therefore indicate an arrangement of orientation-selective voxels beyond what the spatial smoothness of the signal alone would produce. In both monkeys, the early and mid stages contained fewer clusters than expected under the null (monkey R, early: 32 vs. null median 46, p = 0.002; mid: 13 vs. 18, p = 0.039; monkey P, early: 23 vs. 39, p = 0.001; mid: 10 vs. 17, p = 0.010), and these clusters were larger than expected (monkey R, early: 0.0032 vs. 0.0020, p = 0.002; mid: 0.0071 vs. 0.0043, p = 0.010; monkey P, early: 0.0051 vs. 0.0025, p = 0.001; mid: 0.0109 vs. 0.0049, p = 0.001). In the late stage, by contrast, neither property departed from the null (cluster count, monkey R: 18 vs. 15, p = 0.900; monkey P: 13 vs. 13, p = 0.536; mean normalized size, monkey R: 0.0060 vs. 0.0056, p = 0.335; monkey P: 0.0081 vs. 0.0064, p = 0.105). Orientation-selective voxels in the early and mid stages therefore aggregate into fewer and larger clusters than expected by chance, whereas in the late stage this structure is no longer detectable. Although orientation is represented in IT (see previous section), its spatial arrangement there is indistinguishable from that expected under the null. Orientation representation thus follows a systematic spatial organization from early up to mid stages of processing, but becomes less structured and more dispersed in the late stage. This is consistent with orientation no longer being a primary organizing dimension in IT, where mesoscale organization is instead dominated by selectivity for complex visual features.

Whether each stage departs from its own null does not by itself establish that the stages differ from one another. The stages also differ in size and geometry, which could affect cluster count and size independently of orientation organization. We therefore compared the stages directly. We randomly sampled patches from the early and mid stages, matched to the late stage both in volume and in the number of selected voxels, and quantified cluster number and size over 10,000 iterations of this random sampling procedure. Because each sampled patch is matched to the target stage on both quantities that could otherwise drive the difference, this comparison provides the direct test of the transition from mid to late stages. Cluster number in the late stage was significantly higher than in the mid stage but not different from the early stage in both monkeys (monkey R: 18 vs. null median 13 for mid, p = 0.0002, and 15 for early, p = 0.1936; monkey P: 13 vs. 8, p = 0.0070 for mid, and 13 for early, p = 1.0000; Fig. 2E left panels). Mean cluster size (in voxels) was smaller in the late than in the mid stage, reliably so in monkey P but not in monkey R, and did not differ between the late and early stages (monkey R: 9.06 vs. 10.5, p = 0.0616 for mid; 9.06 vs. 9.23, p = 0.8611 for early; monkey P: 9.92 vs. 14.8, p = 0.0260 for mid; 9.92 vs. 9, p = 0.6821 for early). The same matched sampling procedure applied to the early and mid stages, with the mid stage thresholded at its own p < 0.05, showed that clusters in the mid stage were fewer in number (monkey R: 15 vs. 28, p = 0.0002; monkey P: 10 vs. 18, p = 0.0126) and larger in size (22.6 vs. 10.9, p = 0.0002; 19.7 vs. 10.7, p = 0.0102) than those in the early stage (Fig. 2E right panels).

Together, these results reveal a hitherto unexplored property of orientation organization along the ventral stream. From early to mid stages, orientation-selective voxels converge into fewer and larger clusters, aggregating more than expected by chance. In IT, this structure is lost: clusters become more numerous and smaller again, with both properties falling between early and mid stage values, and their arrangement no longer departs from chance. Across stages, the spatial organization of orientation information is therefore broadly consistent with an hourglass shape (Fig. 2F). This implies that orientation follows the mesoscale clustering organization of the visual system up to the intermediate stages V3 and V4, but not beyond them.

### The link between orientation clusters along the hierarchy

So far, we have identified the orientation processing network across the entire visual ventral stream, characterized its distribution in representational space, and examined its clustering organization in cortical space. Next, we ask how the components of this network work together to represent and transform orientation. For this purpose, we explore (1) how the constituent parts of this network are connected, and (2) how information is transformed between these parts.

In order to address the first point, we turned to functional connectivity analysis: we measured Pearson correlation coefficients between pairs of voxels across the three processing stages: early, mid, and late. Since our primary interest was in connectivity, rather than stimulus-driven coactivation, we focused on baseline periods during which no stimulus was presented but the monkey fixated on a blank screen. Deriving connectivity from the inter-block intervals of a task design in this way is an established approach (Fair et al., 2007) and has previously been used to reveal feature- and category-specific coupling within the ventral visual stream that is not attributable to stimulus-evoked coactivation (Norman-Haignere et al., 2012). To minimize the potential influence of residual stimulus-related activity in this analysis, we allowed for 10s of response delay by excluding the initial 10s following stimulus presentation offset (note that results were robust to the precise choice of the length of the delay; see Methods for details). We further removed slow signal drifts by detrending each run and each retained blank block, and scaled each voxel by its residual standard deviation before concatenating blocks across runs, so that the resulting correlations reflect covariation within the blank periods rather than differences in signal level or drift between them.

To relate connectivity to orientation preference, we defined four possible conditions differing in the feature-specificity of connectivity between area pairs: (1) connectivity between top informative voxels (as defined by the absolute t-statistic for the contrast of the two orientations) in upstream and downstream areas (top2top), (2) connectivity between all voxels in the upstream region and the top informative voxels in the downstream region (downtop), (3) connectivity between top informative voxels in the upstream region and all voxels in the downstream region (uptop), and (4) connectivity between all voxels across upstream and downstream regions (Fig. 3A). Then we focused on connectivity trends across these conditions (from top2top to all).

**Fig. 3.**
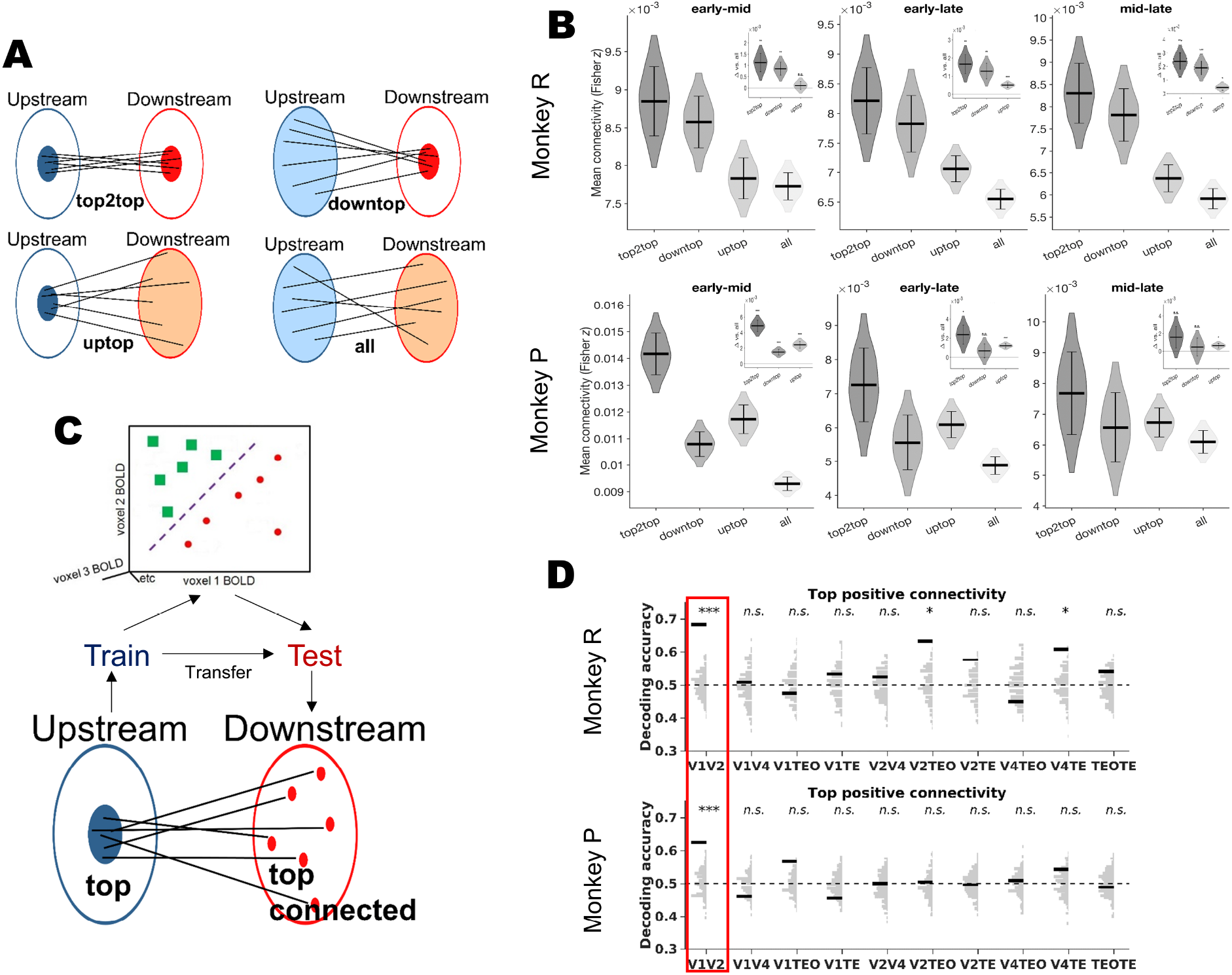
Connectivity and information transfer across the orientation processing network. **A. Top** voxels were defined in each region based on significant orientation-selective contrast t-statistics (*p < 0.05*). Four connectivity types were computed across upstream–downstream region pairs: (1) between top voxels in both regions (**top2top**), (2) between all upstream voxels and top downstream voxels (**downtop**), (3) between top upstream voxels and all downstream voxels (**uptop**), and (4) between all voxels (**all**). **B.** Mean functional connectivity (Fisher-z Pearson correlation) for each connectivity type across the three stage pairs, in both monkeys. Violins show the distribution of the mean across 10,000 bootstrap resamples in which upstream and downstream voxels were resampled with replacement, truncated at the 2.5th and 97.5th percentiles for display. Horizontal bar = observed mean, error bars = bootstrap SEM. Insets: each condition minus the all condition, with two-sided bootstrap p (*** p < 0.001, ** p < 0.01, * p < 0.05, n.s. not significant). **C.** To assess representational transfer, **top** voxels in upstream regions were mapped one-to-one to their strongest connected voxels in downstream regions (**top-connected** voxels; see Methods). A linear SVM classifier was trained on multivoxel patterns from upstream top voxels and tested on the corresponding downstream top-connected voxels. **D.** Decoding accuracy is shown for all upstream–downstream pairs. Thick black bars indicate observed transfer accuracies, and light gray bars represent null distributions generated by random size-matched voxel sampling for statistical comparison. The dashed black line marks chance level (0.5). Asterisks indicate statistical significance: *** denotes *p < 0.001*, ** denotes *p < 0.01*, * denotes *p < 0.05*; and *n.s.* denotes not-significant. The V1–V2 transfer, which was the only pair that proved significant in both monkeys, is highlighted with a red rectangle.

Because each voxel contributes to multiple pairwise connectivity estimates, these estimates share voxel time series and cannot be assumed to be independent. We therefore estimated uncertainty in mean connectivity using a voxel-level bootstrap. In each of 10,000 iterations, upstream and downstream voxels were independently resampled with replacement, and mean connectivity for each condition was recomputed across the pairs formed by the two resampled voxel sets (see Methods).

This analysis revealed a graded pattern of connectivity in all three stage pairs of both monkeys (Fig. 3B): connectivity was highest when orientation-selective voxels were selected at both ends of a connection (top2top), intermediate when selected at one end only (downtop, uptop), and lowest with no selection (all). All eighteen contrasts between a selected condition and the whole-region condition were positive, and fourteen reached significance (Fig. 3B, insets). Top2top was the highest of the three selected conditions in every stage pair of both animals, and exceeded the whole-region condition in five of six comparisons (monkey R: Δ = 0.0011, p = 0.0070 for early-mid; Δ = 0.0017, p = 0.0012 for early-late; Δ = 0.0024, p < 0.001 for mid- late; monkey P: Δ = 0.0049, p < 0.001; Δ = 0.0024, p = 0.0162; Δ = 0.0016, p = 0.2032, respectively), with the sixth in the same direction. The pattern in which top2top is the maximum and the whole-region condition the minimum was preserved in 67% to 100% of bootstrap resamples across the six combinations (median 88%). This suggests (1) a strong anatomical and/or functional route linking orientation processing subnetworks across the hierarchy, and (2) that connectivity is graded by orientation selectivity, increasing with the number of ends at which orientation-selective voxels are selected. We note that the absolute differences in correlation are small, as would be expected from voxel-wise correlations; what the bootstrap establishes is the direction and reliability of these contrasts, not the magnitude of any individual coupling.

One potential concern is that our functional connectivity measure, based on correlation values, might be biased by the response dynamic range of individual voxels. Voxels with higher variance or signal amplitude despite normalization may appear more strongly connected, regardless of whether they carry functionally meaningful orientation information. To test whether this potential bias could explain the observed connectivity pattern, we performed a control analysis using brain regions not involved in orientation processing (MT, LIP, lPFC). If higher connectivity among top informative voxels were simply due to elevated response dynamics, we would expect a similar pattern in these control areas. Specifically, we compared the connectivity of voxels classified as “significant” (p < 0.05) versus “non-significant” (p > 0.05) based on orientation information across all visual-to-control region pairs, matching sample sizes through 1,000 permutations.

We found that functional connectivity between most visual-to-control area pairs was not statistically significant, whereas visual-visual area pairs showed significant connectivity in nearly all cases (Supplementary Fig. 1). These results alleviate concerns about bias in our connectivity measure and further support the observation of strong functional connectivity between voxels representing orientation information.

Overall, these findings demonstrate robust connectivity between clusters representing orientation across the entire ventral processing hierarchy. This connectivity may establish a subnetwork of brain areas that collectively participate in orientation processing. Notably, these connections were observed during blank screen periods, with no active orientation input, indicating that the coupling between orientation-selective voxels is a stable property of the network rather than a consequence of shared stimulus drive. Such stimulus-independent coupling is thought to reflect the underlying anatomical and polysynaptic connectivity of a network, and may have been shaped over time by long-lasting plasticity mechanisms.

### Transformation of orientation information between stages of processing

So far, we have determined orientation-representing clusters along the visual hierarchy as well as the functional connectivity between them. Next, we investigated how information is routed and transformed through those connections. To this end, we grouped visual areas into upstream-downstream pairs. For each pair, we selected a subset of the most orientation informative voxels (defined by absolute univariate contrast t-stats) in the upstream area and identified the corresponding voxels with the strongest connectivity to those in the downstream area. We then trained an SVM classifier in the upstream top informative representation space with a linear kernel and tested its prediction on the corresponding most connected downstream voxels (schematic in Fig. 3C). This allowed us to quantify the extent of shared information between connected upstream and downstream representational spaces. This analysis was conducted across all 15 possible area pairs. To generate a baseline distribution for statistical comparisons, we repeated this procedure using randomly selected, size-matched group of voxels in each downstream area instead of the most strongly connected voxels.

As shown in Fig. 3D, among all pairs tested, only the V1-V2 pair consistently demonstrated above-chance information transfer across both monkeys (decoding transfer = 68.3% in Monkey R and 62.5% in Monkey P; p < 0.001 for both, FDR-corrected for multiple comparisons). Other area pairs either failed to reach statistical significance or showed inconsistent results between the two monkeys. To further validate these findings, we repeated the analysis using voxels with near-zero and top negative connectivity values (see Methods for details). The V1-V2 observed generalization effect did not replicate in these conditions, lending further support to the robustness of the observed V1-V2 information transfer (see Supplementary Fig. 2 for details).

This analysis provides evidence of a direct information transfer from V1 to V2, supporting the existence of communication subspace between these areas. In contrast, we did not observe similar communication between other area pairs. This could indicate either a more complex, non-linear transformation of orientation information across these areas (or a limitation in the precision of the BOLD signal for detecting potential transformations at the microscale).

### Relationship between orientation and object category representation in IT cortex

We found that orientation, as a low-level visual feature, is represented not only in early visual areas but also high-level areas such as IT. This is unexpected according to efficient coding principles: as orientation is already encoded highly precisely in V1, propagation of this information to high-level areas is potentially redundant and hence computationally inefficient. IT cortex, as the apex of visual processing hierarchy, is cardinally involved in representing high-level object categories. If this is the case, then what neural mechanism underlie re- representation of orientation alongside complex object information in IT? And does this re-representation confer any computational benefit to the well-established function of IT, i.e., object representation?

To address these questions, we acquired data using a standard face-object-body (FOB) localizer, presenting blocks of faces, bodies, man-made objects, fruits/vegetables, and phase-scrambled noise in pseudorandom order (Fisher & Freiwald, 2015) (Fig. 4A top panel; see Methods for details). This allowed us to localize face-, body-, and object-preferring regions, respectively, in high-level visual cortex, by contrasting the object categories (Supplementary Fig. 3). Subsequently, we tested whether orientation information is associated with encoding of either of these object categories. To that end, we employed a GLM to predict each voxel’s orientation information (absolute contrast t-stat) in V4, TEO, and TE with its object information (quantified by contrast test comparing object of interest vs. all other object categories) in any of the three categories separately. No significant predictive relationship was found (Supplementary Table 1), suggesting no straightforward link between orientation information and any specific high-level object category in our data.

**Fig. 4.**
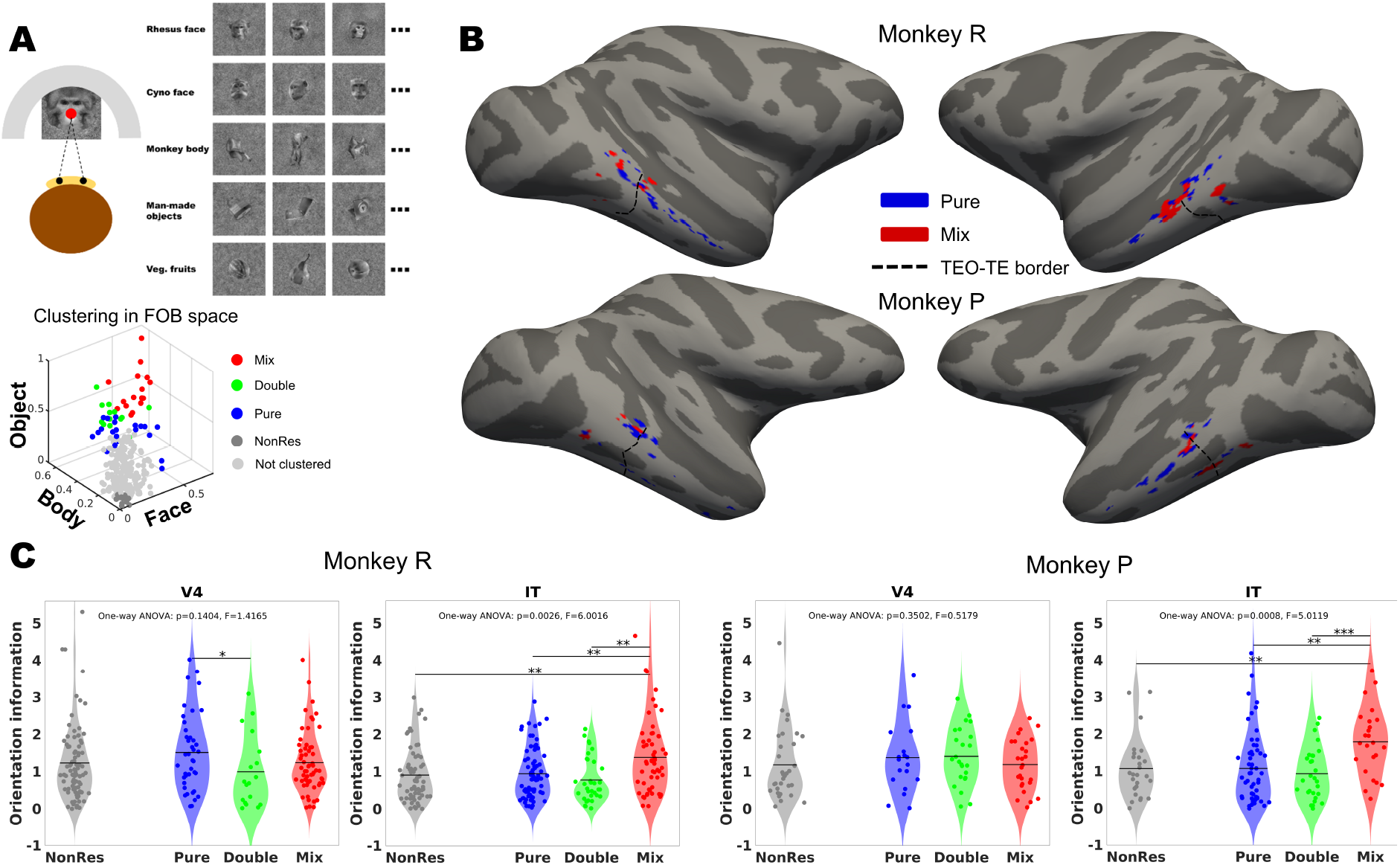
Face–Object–Body (FOB) responsive clusters and their relationship to orientation information. **A.** *Top*: schematic of Face Localizer stimuli and paradigm. *Bottom*: Voxels were classified in FOB space based on their response profile to these three categories. Of particular interest, **Pure** clusters responded to only one category; **Mix** clusters responded to all three categories (faces, bodies, objects). **B.** Pure and Mix clusters projected onto inflated cortical surfaces in both monkeys. Pure clusters appeared more spatially dispersed across IT, while Mix clusters were more localized in posterior IT (TEO and posterior TE). The dashed line marks the TEO–TE boundary. **C.** Violin plots showing the distribution of orientation information (quantified as the absolute value of the orientation contrast t-statistic) across the four voxel types defined in A. Black horizontal lines indicate medians; shaded regions denote standard deviations. One-way ANOVA was used to assess differences across groups and the results appeared on the plots; pairwise comparisons were conducted and only significant results are shown. *** denotes *p < 0.001*, ** denotes *p < 0.01*, * denotes *p < 0.05*.

To further explore the relationship between orientation and high-level object representations, we projected each voxel’s average BOLD response into a 3D representational space defined by face, body, and object dimensions (Fig. 4A bottom panel). We then clustered voxels into four categories: pure (responding to only one category), double (responding to two categories), mix (responding to all three categories), and non-responsive (low response across all categories, used as a control; see Methods for clustering details). Having identified these clusters based on their high-level object response patterns, we compared orientation information across these clusters to determine whether high-level object response patterns relate to orientation encoding. If orientation was linked to object response patterns, two possible scenarios could emerge: either (1) voxels with high selectivity in any category might prefer one orientation due to their object selectivity and/or sparse response properties; e.g., face selective voxels might prefer horizontal orientations while object selective ones preferring vertical/diagonal orientations, or (2) voxels with mixed (multi-variate) responses might represent more orientation information, given their broader responsiveness to diverse features, including orientation.

To appraise these scenarios, we compared absolute orientation information (the absolute contrast t-statistic) across the four identified cluster categories. As shown in Fig. 4C, there was no significant effect of cluster category in V4. Values are reported as mean (standard deviation). For Monkey R, orientation information was 1.2358 (SD 1.0050) in non-responsive, 1.5180 (SD 1.0301) in pure, 1.0012 (SD 0.9615) in double, and 1.2439 (SD 0.8271) in mix-responsive clusters; for Monkey P, values were 1.1825 (SD 0.9465), 1.3764 (SD 0.9364), 1.4103 (SD 0.7665), and 1.1914 (SD 0.6830) in the respective categories (all pairwise comparisons, p > 0.1, FDR-corrected).

In IT, however, we found a significant cluster effect: the mix cluster contained higher orientation information than the other clusters. In Monkey R, orientation information was 0.9153 (SD 0.7207) in non-responsive, 0.9492 (SD 0.6836) in pure, 0.7823 (SD 0.5973) in double, and 1.3889 (SD 0.9826) in mix-responsive clusters. In Monkey P, values were 1.0752 (SD 0.8140), 1.0782 (SD 0.9684), 0.9352 (SD 0.6971), and 1.7941 (SD 0.9103), respectively.

For both monkeys, orientation information in mix-responsive clusters was significantly higher than in non-responsive (monkey R: p = 0.0130, monkey P: p = 0.0060), pure (monkey R: p = 0.0160, monkey P: p = 0.0040), and double (monkey R: p = 0.0130, monkey P: p = 0.0040) clusters (all p-values FDR-corrected). No other pairwise comparisons reached statistical significance (all p > 0.1). These results support the second scenario whereby multi-variate, mix-responsive voxels exhibit stronger orientation representations.

This finding raises a critical question: orientation selectivity or mix-responsiveness, which gives rise to the other? Are these mix-responsive clusters inherently orientation selective, or do they become so because they respond to images containing more coherent, diagonal edge patterns, which is more likely to happen across broader object sets (i.e., a set comprising faces, bodies, objects, etc.) rather than any individual object category? To investigate this, we quantified the orientation content of the images in the different categories in the localizer at both local and global scales (see Methods for details). Statistical comparisons revealed no significant orientation differences between object categories at either level, and no category contained diagonal orientations more than others in our stimulus set (see Supplementary Fig. 4). Thus, the higher orientation information in mix clusters cannot be attributed to an orientation bias in any object category of the localizer set.

This leaves us with another question: Why are multi-variate, mix-responsive voxels more orientation selective? One possibility is that these voxels function as “hubs” in IT cortex, ideally positioned to receive input from earlier processing stages which are strongly tuned for low-level features, and subsequently distribute information throughout IT (Fig. 5A). If this is case, one would expect these hub voxels to exhibit stronger connectivity to upstream visual areas as well as other voxels within IT.

**Fig. 5.**
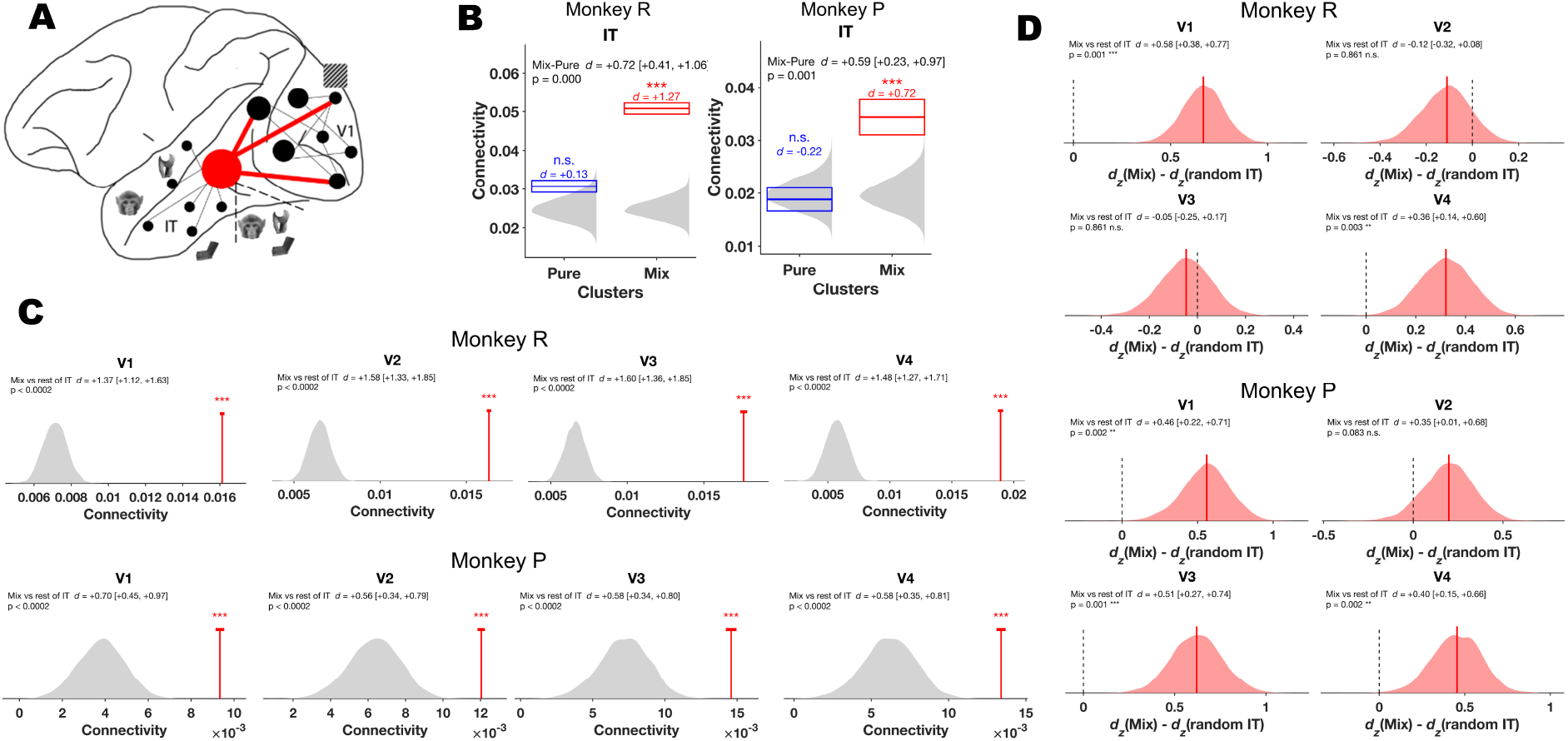
Connectivity of mix IT clusters and the hub network hypothesis. **A.** Schematic of *hub network hypothesis*. Hub voxels are characterized by two properties (1) a mixed category response profile — i.e., responsiveness to multiple object classes — and (2) enhanced functional connectivity both within the same region and to upstream visual areas. **B.** Within-cluster functional connectivity in IT for pure, and mix voxels. Gray violins are the null obtained by shuffling cluster labels across IT voxels (10,000 permutations), which holds cluster size fixed and removes only the assignment. Solid line = observed mean over within-class voxel pairs; open rectangle = 95% confidence interval from a subsampled pair bootstrap (90% of pairs, 5,000 replicates); *d* is Cohen’s *d* computed at the voxel level, comparing each voxels mean connectivity in the class against all remaining IT voxels. **C.** Connectivity of mix clusters to each upstream area (V1 to V4), averaged over all voxels of the target area, compared with the rest of IT. Gray, null distribution of the mean connectivity of size-matched random sets of non-mix IT voxels (5,000 sets); red line, observed mix connectivity, with 95% confidence interval (cap); d, Cohen’s d (mix vs rest of IT, with 95% bootstrap CI). **D.** Preference of mix clusters for orientation-selective upstream voxels (p < 0.05, orientation contrast test). For each IT voxel, preference was defined as its connectivity to orientation-selective voxels minus its mean connectivity to size-matched sets of non-selective voxels from the same area (1,000 draws). Red, distribution of the difference between the preference of mix clusters and that of size-matched random non-mix sets (d_z(Mix) − d_z(random IT), same 5,000 sets as in C); solid line, median; dashed line, zero; d, Cohen’s d of the per-voxel preference (mix vs rest of IT, with 95% bootstrap CI). In C and D, p values are one-sided (mix > rest of IT) and FDR-corrected across the four upstream areas. Significance conventions as in Fig. 2. Mix clusters were more strongly connected than pure clusters within IT, and more strongly than the rest of IT to all upstream areas in both monkeys.

To test this hypothesis, we turned again to connectivity analysis, akin to the one we used to explore connectivity between orientation clusters across the ventral stream. Because cluster categories differ in the number of voxels they contain, we assessed each category against a null distribution obtained by shuffling cluster labels across IT voxels while holding category sizes fixed (10,000 permutations), so that any difference reflects the assignment of voxels to categories rather than their number. Confidence intervals were obtained by resampling voxel pairs, and effect sizes are reported as Cohen’s d computed at the voxel level (see Methods).

Consistent with our prediction, we found that mix IT clusters were more strongly interconnected among themselves than were pure or non-responsive clusters (Fig. 5B). Connectivity values are reported as the mean of z-transformed Pearson correlation coefficients over voxel pairs within each category. In Monkey R, mean connectivity (± 95% confidence interval) was 0.0307 (± 0.0016) for pure, and 0.0508 (± 0.0014) for mix clusters. In Monkey P, values were 0.0188 (± 0.0022), and 0.0344 (± 0.0033), respectively. Mix clusters exceeded the label-shuffle null in both animals (Monkey R: d = +1.27; Monkey P: d = +0.72; both p < 0.001), whereas pure clusters did not depart from it in either (p = 0.061 and 0.717). This elevated connectivity, consistent across both monkeys, provides the first line of evidence for a hub-like property of mix clusters in IT.

We next tested the second hub property: whether this elevated connectivity extends to the upstream areas from which IT receives input. For each upstream area (V1 to V4), we computed the mean connectivity of mix clusters to all voxels of that area. We then drew a random set of non-mix IT voxels, matched in size to the mix clusters, and computed the same connectivity. Repeating this 5,000 times yielded a null distribution of non-mix IT connectivity (Fig. 5C). A mix value within this distribution would indicate no connectivity advantage, whereas a value exceeding it indicates stronger coupling with upstream areas than the rest of IT. Mix clusters exceeded the null in all four upstream areas in both monkeys (all p < 0.001, one-sided, FDR-corrected), with large effect sizes relative to the rest of IT (Cohen’s d [95% CI]; Monkey R: V1 +1.37 [1.12, 1.63], V2 +1.58 [1.33, 1.85], V3 +1.60 [1.36, 1.85], V4 +1.48 [1.27, 1.71]; Monkey P: V1 +0.70 [0.45, 0.97], V2 +0.56 [0.34, 0.79], V3 +0.58 [0.34, 0.80], V4 +0.58 [0.35, 0.81]).

Together, these results show that mix IT clusters exhibit hub-like properties, evidenced by enhanced connectivity both within IT and to upstream visual areas. Since mix clusters also carry more orientation information (Fig. 4C), we next asked whether this is associated with preferential connectivity with highly orientation-selective voxels in other visual areas, or with generally enhanced connectivity with areas more directly involved in orientation processing.

To address this question, we applied the same comparison against size-matched random non-mix IT voxels, but this time to the preference for orientation-selective upstream voxels (Fig. 5D). For each IT voxel, we computed its connectivity to orientation-selective voxels of a given upstream area (p < 0.05, defined by contrast test) minus its connectivity to size-matched sets of non-informative voxels (p > 0.05) from the same area, averaged over 1,000 draws. We summarized this preference as d_z for the mix clusters and for each random non-mix set and formed the distribution of d_z(mix) − d_z(random non-mix). A positive shift of this distribution indicates that mix clusters prefer orientation-selective upstream voxels more strongly than the rest of IT. In Monkey R, the shift was significant in V1 (d = +0.58 [0.38, 0.77], p = 0.001) and V4 (d = +0.36 [0.14, 0.60], p = 0.003), but not in V2 or V3 (both p = 0.861). In Monkey P, it was significant in V1 (d = +0.46 [0.22, 0.71], p = 0.002), V3 (d = +0.51 [0.27, 0.74], p = 0.001) and V4 (d = +0.40 [0.15, 0.66], p = 0.002), but not in V2 (p = 0.083; one-sided, FDR-corrected; d denotes Cohen’s d of the per-voxel preference, mix vs rest of IT). The preference was thus consistent across both animals in V1, where orientation selectivity first emerges, and in V4, the upstream area closest to IT, suggesting that these areas contribute most to the orientation information carried by mix clusters.

The comparisons above show that mix clusters are more strongly connected than the remaining IT population. A more specific question is whether they also exceed narrowly category-selective populations. We therefore compared mix clusters directly with pure clusters (Supplementary Fig. 5). Mix clusters showed stronger connectivity to upstream areas than pure clusters, significantly in Monkey R and as a consistent trend in Monkey P, and a slight preference for orientation-selective upstream voxels (albeit less pronounced than in the comparison with the remaining IT population; Supplementary Fig. 5A, B).

Collectively, these findings identify a subpopulation of IT voxels distinguished by mixed category responsiveness and by markedly elevated connectivity, both among themselves and with upstream visual areas. This elevated connectivity targets upstream areas broadly, but with preference for orientation-informative voxels especially areas V1 and V4. We refer to these mix IT clusters as “hubs”, a term descriptively denoting elevated coupling. These hubs may receive input from upstream processors, support high-level generalized object representation, and distribute this information to other, more specialized clusters throughout IT for further processing.

### Deep convolutional neural networks (DCNNs) exhibit IT-like orientation-object relationship

So far, we have demonstrated the existence of orientation representations in IT and shown how these are linked to heightened connectivity and multivariate response properties, together forming a functional hub at the culmination of the visual ventral stream. We next asked whether deep convolutional neural networks (DCNNs), which share a hierarchical, feedforward architecture with the primate visual system, exhibit a comparable property.

DCNNs trained on core object recognition tasks have been particularly successful in reproducing both primate object recognition behavior and neural response patterns (Cadieu et al., 2014; Yamins et al., 2014). Among the many architectures introduced in recent years, some display stronger brain-like characteristics, as quantified by the Brain-Score metric (Schrimpf et al., 2020), which evaluates model–brain similarity across behavioral and neural benchmarks. We leveraged this metric to directly compare the biological primate visual system with its artificial counterparts.

We ranked the models in descending order of Brain-Score (see Table 1 in (Schrimpf et al., 2020)) and selected those with scores exceeding 0.5, indicating moderate to strong alignment with neural and behavioral data from primate visual system (i.e., explaining over 50% of the variance). We then imported the pretrained versions of these models (trained on the ImageNet dataset) and extracted their penultimate layer responses to the same face–body–object image set used in the fMRI face localizer (see Methods), along with their responses to the two oriented grating stimuli.

We focused on the penultimate layer to enable a direct comparison to monkey IT cortex, as both considered the final stage of visual processing in artificial and biological networks (Cadieu et al., 2014). Using the same clustering procedure that we had applied to the fMRI responses, we identified Pure, Double, and Mix clusters based on their object category response profile, and computed orientation information as the absolute difference in unit responses to the two oriented gratings. We then averaged these values within each model to obtain one value per cluster type and model (Fig. 6B). Across models, mean orientation information (± SD) was 0.140 (± 0.127) for pure, 0.158 (± 0.130) for double, and 0.177 (± 0.141) for mix clusters.

**Fig. 6.**
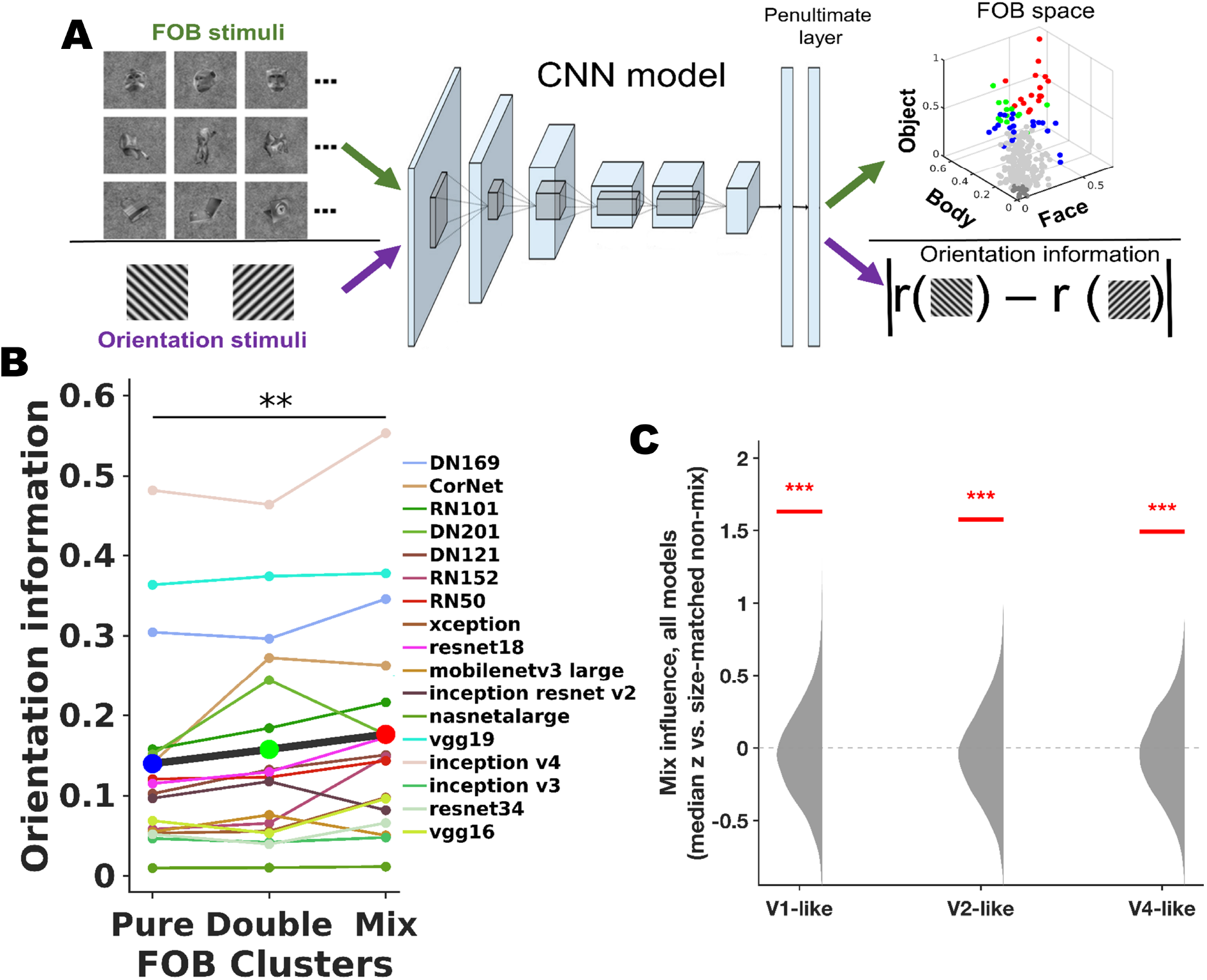
Brain-like DCNNs exhibit similar orientation response patterns across Pure and Mix clusters. **A.** Images of faces, bodies, objects, and two diagonal gratings were input to a set of deep convolutional neural networks (DCNNs) selected based on high Brain-Score values (> 0.5). A clustering approach analogous to that used for IT voxels was applied to the penultimate layer of each model to define **Pure**, **Double**, and **Mix** unit clusters based on category selectivity. Orientation information was computed as the absolute difference in unit responses to the two grating orientations. **B.** Orientation information across the three cluster types for each model. Thin colored lines and circles represent individual DCNN trends; the thick dark-gray line indicates the average trend across all models. Solid, slightly enlarged red, green, and blue circles mark the mean orientation information across models for Mix, Double, and Pure clusters, respectively. Only the Pure vs. Mix comparison reached statistical significance (Wilcoxon signed-rank test across models, FDR-corrected; ** *p < 0.01*). DN: DenseNet; RN: ResNet. **C.** Upstream influence of mix units in the penultimate layer, aggregated across the 17 DCNNs, for V1-, V2-, and V4-like layers. For each architecture, gradient-based sensitivity analysis quantified how strongly activations in each upstream layer contributed to the response of each penultimate unit (see Methods); units with zero influence were excluded. For each model and layer, the mean influence of mix units was compared with a null distribution of size-matched random sets of non-mix units (10,000 sets) and expressed as z relative to that null. Gray, null distribution of the median z across models, obtained by taking the median of the models’ standardized null values for each of the 10,000 sets; red line, observed median z across models. P values are two-sided permutation p values, FDR-corrected across the three layers. Across models, mix units received higher upstream influence than size-matched non-mix units from all three layers, paralleling the elevated upstream connectivity of IT hub voxels in the fMRI data (Fig. 5C).** denotes *p < .01,* *** denotes *p < .001*.

Pairwise comparisons of clusters across network architectures revealed that orientation information was significantly higher in mix compared to pure clusters (mix vs. pure: p = 0.0042), while the differences between mix and double clusters (p = 0.0759) and between pure and double clusters (p = 0.0759) did not reach significance (sign-rank test, FDR-corrected). These findings replicate the effect observed in IT cortex across both monkeys. It bolsters the observed link between orientation representation and object category response profile at the final processing stage and shows that this relationship extends to brain-like artificial networks.

Finally, to test whether a functional hub resembling IT hub voxels also emerges in the penultimate layer of DCNNs, we quantified the influence of upstream layers (V1-, V2-, and V4-like) on penultimate units (see Methods). Following the same logic as in the fMRI data, we compared, for each architecture, the mean upstream influence of mix units with a null distribution of size-matched random sets of non-mix units, and combined the resulting effects across models. Across the 17 models, mix units received higher upstream influence than size-matched non-mix units from all three upstream layers (median z = 1.63, 1.58, and 1.50 for V1-, V2-, and V4-like layers, respectively; all p < 0.001; Fig. 6C), with positive effects in 12 of 17 models. Notably, this pattern closely mirrors the elevated upstream connectivity of IT hub voxels in the fMRI data (Fig. 5C). As in the monkey data, a direct comparison with narrowly category-selective pure units showed the same pattern across models (Supplementary Fig. 5C).

Together, these findings point to a shared organizational principle between the primate visual system and DCNNs: both converge on forming a hub in the final stage of processing that integrates and transforms information from earlier layers. This parallel suggests that orientation-linked hubs may constitute a general computational strategy for supporting flexible object representations.

## Discussion

In this study, we systematically explored the network processing orientation information across the entire macaque visual ventral stream hierarchy. Our results demonstrate that orientation information is decodable, with varying accuracies, from multi-voxel pattern responses across areas, ranging from early (V1, V2) to late stages (IT) of processing. Spatially, orientation information was organized in an intriguing hourglass shape across the visual hierarchy: distributed representations in V1 and V2 converged into dense clusters in V3 and V4, while becoming again more numerous and scattered in IT. In contrast, in representational space, we found a steadily increasing distribution of information along the visual hierarchy. This was evidenced by a growing distance between univariate and multivariate information metrics (described as sigProp and peakProp in Results section) (Fig. 2B) comparing V1 and IT, and this difference persisted when V1 was subsampled to match IT in both size and univariate selectivity. Orientation-responsive voxels at the highest stages of processing in IT did not show selectivity for complex objects, a property shared with orientation-responsive units in top layers of DCNNs. Orientation-responsive voxels were also strongly interconnected within IT as well as more variably to earlier visual areas. Taken together, this suggests that orientation information is redundantly encoded across the ventral stream but may be used for different computational roles, requiring different representational formats and spatial organization, at different stages of processing.

### Organization of orientation information along the ventral stream

The hourglass-shaped pattern of spatial clustering that we found along the ventral stream is notable for (at least) two reasons: (1) it deviates from the typical convergence logic of the ventral stream (from early to late stages) (Y. Liu et al., 2020; Vanduffel et al., 2002), and (2) it contrasts with other low-level features like color (Lafer-Sousa & Conway, 2013) or high-level features like faces, bodies, and objects (Moeller et al., 2008; Pinsk et al., 2009; Tsao et al., 2006; Zhu et al., 2025), which exhibit a clustered and patch-like organization at the highest stages of processing (Supplementary Fig. 3). Interestingly, accounts of the structure of the ventral stream based on graph-theory (rather than on the laminar pattern of connectivity) allocate V4, where we observed the largest degree of spatial clustering for orientation, to the top of the hierarchy, whereas V1 and anterior IT rank on a similar, lower level, thus mirroring the hourglass shape we observed (Sporns, 2016; Sporns et al., 2007).

Spatial clustering in the ventral stream is thought to arise from distance-dependent constraints on neural computation (Blauch et al., 2022; Z. Lu et al., 2025; Margalit et al., 2024) and may be critical, e.g., for local computations such as sharpening within stimulus domains (Sompolinsky & Shapley, 1997). How and why such computations would differ between areas for the same feature may depend on the specific tasks to which the respective areas contribute (Ahissar & Hochstein, 2004; Fang et al., 2022; Manenti et al., 2023).

Our results add a further observation relevant to these constraints: the increase in cluster number from mid to late stages was accompanied by a rise in the number of isolated, unclustered voxels, without a proportional reduction in the size of the clusters that formed. Orientation selectivity in IT thus appears fragmented rather than merely rescaled, which may reflect a relaxation of the spatial constraints that shape clustering at earlier stages, where representations are more directly tied to retinotopic organization.

The increasingly distributed nature of orientation information from V1 to IT in representational space further suggests that readout strategies may differ across the ventral stream: in early stages, orientation representation may be more focused on a select subset of high-informative voxels, consistent with selective readout (Nadler & DeAngelis, 2005; Purushothaman & Bradley, 2005). As processing progresses, orientation representations may become less dependent on these highly informative voxels and increasingly distributed across less informative voxels in later stages, consistent with a more global readout model (Shadlen et al., 1996).

An alternative account would attribute this trend to lower signal quality of anterior temporal cortex, where fewer voxels reach univariate significance and multivoxel decoding must therefore recruit a larger proportion of the area. Our matched sampling analysis argues against this: when V1 was subsampled to match IT not only in size but in its full distribution of univariate selectivity, the observed IT distance still exceeded the resulting null. Since this matching equates the proportion of individually significant voxels by construction, the difference must arise from how multivoxel information accumulates across voxels rather than from how many voxels are individually informative.

### Orientation transformation along the visual ventral stream

In addition to identifying the organization of orientation clusters across the visual hierarchy, we investigated how these clusters are interconnected and how orientation information transforms between them. Significantly informative orientation voxels exhibited stronger connectivity with one another compared to non-informative voxels. This selectively enhanced connectivity was evident during the baseline periods, in which no stimulus was presented. Coupling measured during such intervals has been shown to carry feature- and category-specific structure in ventral visual cortex that cannot be attributed to stimulus-evoked coactivation (Al-Aidroos et al., 2012; Norman-Haignere et al., 2012) and we take our findings to reflect (polysynaptic) pathways within and between brain areas forming an orientation processing subnetwork in the ventral stream. This resembles what has been reported on the basis of anatomical connections for other visual features such as color: For example, color-biased blobs in V1 are specifically and reciprocally connected with color-biased thin stripes in area V2 (Federer et al., 2021; Livingstone & Hubel, 1984). We note, however, that our measure is correlational and cannot establish the directness of these pathways.

Within this network, our findings reveal a direct linear interaction for orientation information between areas V1 and V2. This aligns with an electrophysiological study that identified a communication subspace linking subsets of V1 and V2 neurons in anesthetized macaques viewing oriented gratings (Semedo et al., 2019). Among all other visual area pairs we analyzed, we did not observe similar effects, suggesting privileged communication between V1 and V2. This may arise from the particularly strong structural feedforward and feedback connectivity between V1 and V2 that exceeds that of other areas (H. Li et al., 2024). Additionally or alternatively, the absence of a linear interaction between the other area pairs could be indicative of more complex non-linear transformations between these regions (and/or a limitations of fMRI in capturing fine-grained circuit dynamics).

### Orientation information in IT cortex

The ventral visual stream has long been considered an epitome of hierarchical processing systems, wherein simple visual features (e.g. orientation, color, spatial frequency) are extracted early, and progressively integrated into more complex representations downstream (Kravitz et al., 2013; Pollen et al., 1971). In this framework, simple features are not redundantly represented in higher visual areas like IT. However, lesions of higher visual areas such as V4 (Weerd et al., 1996) and even TE (Dean, 1978) can induce deficits in orientation discrimination, and a small number of neurophysiological studies reported responses of IT neurons to oriented gratings or bars (Gross et al., 1972; Maunsell et al., 1991; Orban & Vogels, 1998; Pollen et al., 1971, 1984; Vogels & Orban, 1993). Such effects have previously been attributed to task-demands: e.g., long time constants of neural activity in higher-level cortex are thought to support temporally successive orientation discrimination (Orban & Vogels, 1998; Vogels et al., 1997). Yet, our task did not require discrimination nor did it pose any memory demands.

An alternative hypothesis states that responses to oriented stimuli in high-level cortex arise as a byproduct of responses to complex objects. This hypothesis is supported by a number of recent fMRI studies investigating the dependence of high-level feature representations on low-level attributes in humans (Fan et al., 2021; Henderson et al., 2023; Nasr & Tootell, 2012; Rice et al., 2014) and non-human primates (Bao et al., 2020; Kumar et al., 2019; Yao et al., 2023). They find that orientation content of images can predict, to varying degrees, each voxel’s BOLD response to those images. For example, face- and body-selective areas show a bias toward diagonal orientations, while scene-selective areas favor vertical and horizontal orientations. Another possibility is that orientation itself is not explicitly represented in IT but emerges as an epiphenomenon of high-level visual features or object category representations, which are inherently rich in orientation content (Henderson et al., 2023). However, all above-mentioned neuroimaging studies measured IT’s orientation response indirectly by analyzing the orientation content of complex objects, often using Gabor filters. This approach has two critical limitations (i) complex images typically contain abundant orientations, making it difficult to isolate responses to specific ones, and (ii) IT responses to such images may result from non-linear interactions between orientations, which linear separation methods cannot fully capture. Given these concerns, we adopted a more direct measurement of orientation responses using classical sine-wave grating stimuli. If orientation representation in IT is merely a byproduct of complex object processing, it should vanish when simple gratings are presented. However, we found that approximately 13% of IT voxels exhibited significant orientation responses. This makes it unlikely that IT’s orientation response is purely a byproduct of object processing (Supplementary Fig. 3). While we cannot entirely rule out the hypotheses of task-specificity or complex object dependence for orientation representations in IT, we put forward and evaluate an alternative hypothesis, which we elaborate on in the following sections.

### Orientation information in IT cortex: The Hub Hypothesis

To delve deeper into the repeated representation of orientation along the ventral stream, we explored the relationship between orientation and complex objects. Interestingly, we found that voxels responding to multiple object categories, referred to here as mixed responsive voxels, were more diagnostic to orientation than strongly category-selective voxels (Fig. 4C). That voxels responsive to objects do not overlap with voxels responding to the constituent features bears some resemblance to the observation that V4 clusters tuned to curves and corners do not overlap with clusters tuned for orientation (Jiang et al., 2021). Mixed responsive voxels in our data were primarily located in IT, distributed along cortex rather than clustered, and tended to localize more posteriorly, near or within TEO.

Moreover, these mixed responsive voxels exhibited stronger connectivity among themselves within IT, an effect that was robust in both animals. They were also more strongly connected to upstream areas V1 and V4. Known anatomical pathways linking IT with earlier visual areas include reciprocal connections between TEO and V4 (Distler et al., 1993) and direct feedback projections from TEO and TE to V1 (Rockland & Van Hoesen, 1994). These pathways provide a potential anatomical substrate for the observed coupling, although our functional connectivity measure cannot determine its direction or distinguish direct from polysynaptic interactions. Together, our observations identify an IT subpopulation characterized by elevated connectivity and by a diverse response profile encompassing both orientation and multiple object categories. We refer to these voxels as “hubs”. Due to their high connectivity and mixed responsiveness, these hubs could play a critical role in various functions:

1. By virtue of their mixed responsivity (Johnston et al., 2020; Johnston & Freedman, 2023; Rigotti et al., 2013), hubs could make decision making (Johnston et al., 2020; Kira et al., 2023; Zhou et al., 2021) and learning processes (Barak et al., 2013; Karami & Schwiedrzik, 2024) more flexible. Indeed, higher-order areas such as IT cortex have been suggested to be more plastic than lower areas (Ahissar & Hochstein, 2004).
2. Given the relative lack of specialization and strong connectivity with other IT and upstream neurons, hubs are ideally positioned to be modulated by downstream regions to flexibly render the information required for a task at hand (W. Li, 2016; Zylberberg, 2018). This capacity makes them conducive to contribute to adaptive behavior and one-shot learning (Donoghue et al., 2023; Kondat et al., 2024; Lee et al., 2015; Ramirez-Cardenas et al., 2016; Sorscher et al., 2022).
3. Because of their broad upstream connectivity, hubs in IT could take privileged roles in processes that require integrating information across earlier areas, such as directing attention. Indeed, recent studies indicate that area TEO, where we found most hubs, may play a key role in attentional deployment, not only for complex objects but also for low-level features (Stemmann & Freiwald, 2019).

### Hub units also emerge in Deep convolutional neural networks (DCNNs)

Deep convolutional neural networks (DCNNs) have been remarkably successful in explaining a plethora of phenomena in primate vision, from object recognition to the neural response pattern similarities. In the present study, we reveal an unexplored property of the primate visual system: the emergence of a functional hub in the final processing stage, characterized by multivariate responsiveness and enhanced connectivity to upstream regions. Motivated by the success of DCNNs as models of visual processing, we asked whether a similar hub-like property emerges in their architecture, particularly the ones with higher brain similarities. Indeed, we found that DCNNs also encode orientation information in their penultimate layer, where it is linked to multivariate responsiveness. Furthermore, across models, mix units showed greater upstream influence than size-matched sets of non-mix units, with a similar pattern in direct comparisons with pure units. This pattern parallels the upstream connectivity difference we observed for hub voxels in macaque IT cortex. Our findings suggest that narrow category selectivity is not the only organizing principle of high-level representations in networks optimized for object recognition. Mix-responsive units combine greater orientation information with enhanced upstream influence, linking sensitivity to a basic visual feature to a distinctive position within the processing hierarchy. This extends the correspondence between DCNNs and IT beyond category-selective representations to the relationship between response diversity and network organization.

### Limitations

In this study, we employed full-field sine-wave gratings at a single spatial frequency of 2 cpd. This choice was motivated by the fact that 2 cpd falls near the median preferred spatial frequency of macaque V1 neurons (Bredfeldt & Ringach, 2002; De Valois et al., 1982) and within the range of peak perceptual sensitivity for natural images (Bex & Makous, 2002), making it well suited for capturing orientation responsiveness across the entire visual cortex. However, we acknowledge that spatial frequency tuning exists throughout the visual hierarchy (M.-L. Liu et al., 2024; Toosi et al., 2025), and therefore our protocol may have missed voxels with narrowband sensitivity to substantially lower or higher spatial frequencies. Additionally, the use of full-field gratings may not have optimally driven neurons with strong surround suppression or end-stopping, potentially leaving such populations undetected by our present protocol. Finally, we used conventional BOLD imaging without a contrast agent such as Magnetic Iron Oxide Nanoparticles (MION). While contrast-enhanced imaging can improve sensitivity and spatial specificity in macaque fMRI, our BOLD protocol was sufficient to reliably recapitulate well-known polar angle and eccentricity maps across early visual cortex as well as face patches in IT cortex (Supplementary Figs. 3 and 5), confirming that it captures the functional organization relevant to the present study.

### Conclusion

In conclusion, our findings suggest that low-level representations in high-level areas like IT emerge as a result of network connectivity and may serve novel computational roles beyond IT’s core function of object processing. This phenomenon likely represents an adaptive feature of the visual ventral stream, enabling additional functions such as learning, flexibility, and attention, rather than being a byproduct of high-level visual processing itself. Our fMRI and DCNN data suggest the existence of “Hub” neurons in macaque IT, characterized by mixed selectivity to diverse object categories as well as low-level features. Future research should focus on confirming the existence of these neurons in IT and further exploring their computational roles. This includes their potential contributions to high-level visual processing and higher-order cognitive functions such as learning, memory, decision-making, attention, and adaptive behavior in general.

## Acknowledgements

We would like to thank Wim Vanduffel and Arjen Alink for input on the project; Igor Kagan for support; Ronja Mielsch, Daniela Lazzarini, Leonore Burchardt, and Sina Plümer for help with animal training and care; and Tamara Becker, Olga Batura, and Annette Schrod for veterinary care. This project has received funding from the European Research Council (ERC) under the European Union’s Horizon 2020 research and innovation programme (Grant agreement No. 802482) and from the German Research Foundation’s Emmy Noether Program (SCHW1683/2-1). This work used the Scientific Compute Cluster at GWDG, the joint data center of Max Planck Society for the Advancement of Science (MPG) and University of Göttingen. This research was supported in part by the Intramural Research Program of the National Institutes of Health (NIH). The contributions of the NIH author(s) are considered Works of the United States Government. The findings and conclusions presented in this paper are those of the author(s) and do not necessarily reflect the views of the NIH or the U.S. Department of Health and Human Services. The content of this work is solely the responsibility of the authors and does not necessarily represent the official views of the European Union, or the European Research Council. Neither the European Union nor the granting authority can be held responsible for them.

## Author contributions

B.K., Conceptualization, Methodology, Investigation, Formal Analysis, Visualization, Data curation, Writing – original draft preparation; T.N., Methodology, Investigation, Data curation; S.S.P., Methodology, Investigation, Data curation; C.M.S., Conceptualization, Methodology, Writing – original draft preparation, Supervision, Project administration, Funding acquisition.

## Materials and methods

### Subjects and procedures

We acquired functional MRI data from two adult male macaque monkeys (*Macaca mulatta;* monkey R, 10-11 kg, 8 yrs, and monkey P, 6-7 kg, 7 yrs). No power calculation was performed, but sample size was chosen to be in accordance with previous studies (Moeller et al., 2008; Nigam & Schwiedrzik, 2024). The animals were housed in pairs or social groups at the German Primate Center (DPZ) in accordance with German and European regulations. The facility provided a stimulating environment for the animals, including various toys and structures, natural and artificial lighting, and access to outdoor spaces, surpassing the space requirements outlined in European regulations. All procedures were approved by the responsible regional government authority (Niedersächsisches Landesamt für Verbraucherschutz und Lebensmittelsicherheit - LAVES).

Both animals underwent surgery under general anesthesia and sterile conditions to implant an MRI-compatible cranial head-post (PEEK) embedded in bone cement (Palacos, Heraeus Kulzer GmbH), anchored by MR-compatible ceramic screws (Rogue Research) (Dominguez-Vargas et al., 2017). Prior to the main fMRI data acquisition phase, both monkeys completed substantial training for horizontal chair acclimation and passive fixation tasks using positive reinforcement. Head position was stabilized throughout training and data acquisition using the implanted head-posts, allowing for cleaning of the implants, accurate recording of gaze/pupil diameter, and minimizing motion artifacts during MRI experiments.

This fMRI study was conducted in non-human primates for several reasons, including: (1) the extensive existing knowledge of primate visual system enables direct comparison with well-established electrophysiological and neuroimaging findings, in particular concerning orientation representations in early visual areas and object/category representations in higher visual areas; (2) using trained, head-stabilized non-human primates permits prolonged, high-resolution data acquisition; and (3) any observed effect can lay the foundation for immediate testing of hypothesized underlying mechanism using electrophysiological and causal techniques - approaches uniquely feasible in non-human primate models (Milham et al., 2022).

### MRI data acquisition

We used a 3T scanner (MAGNETOM-Prisma, Siemens Healthineers, Erlangen, Germany) at the DPZ to collect both anatomical and functional MRI data. Data acquisition protocols followed community guidelines (Autio et al., 2021). High-resolution anatomical MRI data were acquired using a T1-weighted magnetization-prepared rapid gradient echo (MPRAGE) sequence (field of view [FOV] 128 mm, voxel size = 0.5 × 0.5 × 0.5 mm, repetition time [TR] = 2.7 s, echo time [TE] = 2.96 ms, inversion time [TI] = 850 ms, bandwidth [BW] = 220 Hz/Px, flip angle [FA] = 8 degrees, 240 slices, 11 cm loop coil, Siemens Healthineers). The monkeys were anesthetized during these scans using isoflurane (1.5%–2%) and were positioned in an MR-compatible stereotactic frame (Kopf Instruments). We acquired functional images in awake animals using 8-channel phased-array receive surface coils with a horizontally oriented single loop transmit coil (H. Kolster, Windmiller Kolster Scientific). During these scans, monkeys were seated in sphinx position inside an MR-compatible horizontal chair (Valette et al., 2006). Each functional time series consisted of whole-brain gradient-echo planar images (EPI; FOV = 96 mm, voxel size = 1.2 × 1.2 × 1.2 mm, TR = 2 s, TE = 27 ms, BW = 1302 Hz/Px, echo spacing [ESP] = 0.93 ms, FA = 76 degrees, 43 slices) acquired in interleaved order with two times generalized autocalibrating partially parallel acquisitions (GRAPPA) acceleration.

### Stimuli and task

The paradigm — including stimulus presentation and reward delivery — was programmed and conducted using Psychtoolbox (Brainard & Vision, 1997) running in MATLAB (The Mathworks). Visual stimuli were rear-projected onto a screen positioned 85 cm in front of the monkey inside the scanner, using a video projector (Epson EB-G5600, refresh rate 60 Hz, resolution 1024 × 768 px). Eye position was tracked using a video-based eye tracker (SR Research Eyelink 1000 NHP Long Range Optics) sampling at 1000 Hz.

In all experiments, monkeys were rewarded with fluid for maintaining fixation within a 2° diameter central fixation window for 3.7 ± 0.2 seconds. This fixation duration was chosen to minimize temporal overlap between reward delivery and image acquisitions (TRs). Only runs with fixation performance ≥ 85% were later included in the analyses.

#### Orientation mapping

We employed an fMRI block design to map orientation-selective responses across the entire macaque cortex. To this end, full-field oriented gratings (full contrast; spatial frequency = 2 cyles per degree) were presented to robustly activate all visually responsive regions across the entire visual field. A central red fixation point (0.5° diameter) was continuously displayed throughout each block to guide fixation. Parameters followed previous studies (Alink et al., 2013; Henriksson et al., 2008; Tong & Pratte, 2012; Vanduffel et al., 2002).

In Monkey R, six orientations (0°, 45°, 67.5°,90°,112.5°,135°) were each presented for 32 seconds, interleaved by 24 seconds of blank black background. Even though six orientation angles were presented to this monkey, only the two diagonal angle conditions (45° and 135°) were included in analysis for the present study. In Monkey P, only the two diagonal gratings (45° and 135°) were presented, each for 26 seconds, interleaved with 26 seconds of blank black background. Each orientation condition was repeated seven times.

Gratings flickered at 6 Hz to maximize the evoked BOLD response, and their phase was randomized across four values (0, π/4, π/2, 3π/4) to optimally stimulate both simple and complex cells in early visual cortex.

In total, 60 runs in Monkey R and 15 runs in Monkey P met the inclusion criteria of ≥85% fixation stability to be included in analysis. Over all usable runs, each diagonal grating was presented 60 times in Monkey R and 104 times in Monkey P.

#### Polar angle and eccentricity mapping

For polar angle and eccentricity mapping, we followed established procedures (Kolster et al., 2014) To obtain polar angle maps, we presented visual stimuli consisting of a wedge with an arc length of 30°, rotating either clockwise or anticlockwise against a black background with a red central fixation point of 0.5° diameter. The arc moved at the speed of 6.3/s. The wedge contained full-contrast black-and-white checkerboard patterns composed of 18 concentric circles each containing 40 radial black–white segment pairs. To enhance the evoked response, the black and white elements alternated at a flicker rate of 2.5 Hz. Each run included 5 full rotation cycles in Monkey R, and 8 cycles in Monkey P (57.1428 seconds per cycle), followed by 12 seconds of black background presentation at the end.

For eccentricity mapping, expanding or contracting black-and-white checkerboard pattern rings were presented around a 0.5° red central fixation point. A total of 18 rings were used to map the visual field from 0.4° to 16.5° of eccentricity. Each ring was composed of three concentric circles, except for the two most peripheral rings, which contained two and one circle(s), respectively. Each circle contained 24 angular black–white segment pairs and spanned approximately 0.92 ° of visual field in eccentricity.

During each presentation cycle, rings were presented sequentially sweeping linearly across the visual field— each for 2 seconds in Monkey R (36 s total) and 3.1746 seconds in Monkey P (57.1428 s total). Each run consisted of 8 cycles in Monkey R, and 12 cycles in Monkey P, followed by a 12-second blank, black background.

In total, we acquired 27 polar angle and 17 eccentricity mapping runs for Monkey R, and 14 polar angle and 24 eccentricity runs for Monkey P.

#### Face-Object Body localizer

We used a standard face–object–body (FOB) localizer protocol as described in (Fisher & Freiwald, 2015; Nigam & Schwiedrzik, 2024; Schwiedrzik et al., 2015). Monkeys fixated on a central red fixation dot (0.5° diameter) while viewing grayscale images from the following categories: monkey faces, body parts/headless bodies, man-made objects, fruits, and phase-scrambled images. Each image was presented for 0.4 seconds, and each category block lasted 36 seconds, followed by a 24-second presentation of a blank gray background. In total, 60 runs in Monkey R, and 51 runs in Monkey P passed the inclusion criteria and were used in the analyses. Data from Monkey P were also used in another publication (Nigam & Schwiedrzik, 2024).

### Analyses

#### General preprocessing

Anatomical images were intensity normalized, skull-stripped, and segmented using Freesurfer (v5.3, https://surfer.nmr.mgh.harvard.edu/) (Fischl, 2012). To create inflated cortical surface reconstructions, the gray–white matter boundary in the skullstripped anatomical scans was reconstructed, smoothed, and inflated separately for each hemisphere (Dale et al., 1999; Fischl et al., 1999). Preprocessing of functional data was performed with custom scripts using Freesurfer’s functional analysis stream, FS FAST (Fischl, 2012), AFNI (https://afni.nimh.nih.gov/) (Cox, 1996), and the JIP toolkit (https://www.nitrc.org/projects/jip/). We performed slice-wise motion-correction in each run using AFNI’s 3dAllineate with terms for cubic warping in the phase encoding direction, shifts, rotations, scaling, and skewing. This process was followed by slice-time correction using FS FAST. Geometric distortions of the functional volumes were corrected by mutual information-based non-linear alignment to the high-resolution anatomical scans as implemented in JIP.

#### Visual Region of Interest (ROI) definition

We employed Fourier analysis to obtain spatial selectivity of voxels from polar angle and eccentricity mapping sessions (Arcaro & Livingstone, 2017; Bandettini et al., 1993; Engel et al., 1997). We determined the phase and amplitude of the periodic signal that matched stimulus frequency using Fourier transform of the mean time series accounting for a 4-second hemodynamic lag. The resulting phase angle data were registered and projected onto individualized inflated cortical surfaces for both monkeys (Supplementary Fig. 6).

We followed established procedures to define visual regions (Arcaro et al., 2011; Brewer et al., 2002; Janssens et al., 2014; Van Essen et al., 2001). Boundaries were delineated based on color-coded phase maps where clearly identifiable. In cases where boundaries could not be reliably defined—particularly in higher-order visual areas and associated control regions (MT, LIP, and lateral prefrontal cortex [lPFC])—we used the D99 atlas (Saleem et al., 2021), individually registered for each monkey, to complete the delineation.

#### Univariate analyses

We used a General Linear Model (GLM) in FS FAST to estimate the contribution of each diagonal orientation to the BOLD signal. Prior to the analysis, we convolved the predictors of interest with the canonical hemodynamic response function (HRF) for the blood-oxygenation level dependent (BOLD) signal (Leite et al., 2002). Nuisance regressors included 3D motion, eye position, and reward delivery. To determine the differential effect of orientation, we contrasted the two diagonal orientations (45° vs. 135°) and obtained t-statistics and corresponding p-values. These were used to identify orientation-selective voxels. We use univariate “orientation information” to refer to orientation-dependent differences in the measured BOLD response, indexed by the absolute contrast t-statistic. This terminology follows the use of “information” to describe statistically discriminable responses without requiring an information-theoretic measure (Kriegeskorte et al., 2006; Mur et al., 2009), including previous applications to voxelwise t-statistics (Gotsopoulos et al., 2018). We note that this index can in principle additionally depend on measurement sensitivity and noise.

#### Orientation representations across visual hierarchies; MVPA analysis

To quantify orientation representations in each visual area, we measured decoding accuracy in the voxel patterns using multivoxel pattern analysis (MVPA) as described in (Harrison & Tong, 2009). We began by regressing out nuisance variables (eye position, 3D motion, and reward). Eye position and reward were convolved with the HRF to match the temporal dynamics of the BOLD signal. Only residuals, which retain the main task-related signal while excluding the impact of external confounds, were used for further analysis.

Next, we averaged the residual BOLD time series within each block containing one of the three conditions: blank screen, clockwise grating (45°), or anti-clockwise grating (135°), to obtain the average BOLD activity per condition per run. We discarded the first 4 seconds (2 TRs) of each block to account for the hemodynamic response delay. This yielded a matrix of average BOLD responses for the three conditions across voxels within each region of interest (ROI), forming the basis of multivoxel representational space.

On this space, we trained a linear support vector machine (SVM) classifier to discriminate between the two orientation conditions (clockwise vs. anticlockwise), using two-thirds of the repetitions for training and one-third for testing. We repeated this process on randomized train/test splits for 100 repetitions (cross-validation) and computed the mean and standard deviation of decoding accuracy, as presented in Fig. 1B. To establish a baseline for statistical comparison, we shuffled orientation labels and repeated the decoding analysis 1000 times to generate a null distribution. Each observed decoding accuracy per ROI was then tested against this distribution using a permutation test. This analysis was performed over visual ROIs (V1, V2, V3, V4, TEO, and TE), as well as in control regions MT, LIP, and lateral prefrontal cortex (lPFC). To track the progression of orientation information across the visual hierarchy, we also grouped visual ROIs into three processing stages: early (V1, V2), mid (V3, V4), and late (TEO, TE /IT).

#### Orientation organization in cortical volume space

To quantify how orientation information is spatially organized in the cortical volume, we clustered voxels showing high univariate orientation selectivity, as determined by the contrast t-statistic (see section “Univariate analyses”).

#### Voxel selection

Rather than thresholding each processing stage independently, we fixed the proportion of selected voxels across stages. We first determined the proportion of voxels in the late stage reaching p < 0.05 in the orientation contrast (sigratio). We then applied this same proportion to the early and mid stages by selecting, within each, the voxels with the highest absolute t-statistic until sigratio × N voxels were selected. This ensures that between-stage differences in cluster number and size cannot arise from differences in how many voxels survive selection, which would otherwise vary with signal quality across the brain.

#### Clustering

Selected voxels were grouped using the following criteria: (i) selected voxels immediately adjacent in 3D cortical space were grouped together; (ii) voxels with no immediate neighbor were labeled as isolated and excluded from clusters. Adjacency was defined by 26-connectivity, in which two voxels are neighbors if they share a face, an edge, or a corner. We implemented the clustering using a region-growing algorithm. Beginning with the most posterior-medial selected voxel (arbitrarily chosen), we recursively searched its immediate 3D neighborhood for other selected voxels. If a neighboring voxel was found, it was grouped into the same cluster; otherwise, it was labeled as isolated. This process continued until all voxels were either clustered or labeled as isolated. To maintain spatial cohesiveness and reduce branching, we pruned clusters by reclassifying voxels with fewer than two neighbors as isolated. We recorded, for each stage, the number of clusters, the mean cluster size normalized by the total volume of that stage, and the number of isolated voxels.

#### Null distributions

We assessed the observed cluster properties against two null distributions, each constructed with 1,000 permutations and each holding the number of selected voxels at its observed value.

In each permutation, the two orientation labels were swapped at random for each pair of anticlockwise and clockwise blocks (i.e., the sign of each block-wise difference image was flipped at random; 60 pairs in monkey R, where each block constituted a separate run, and 104 in monkey P), the paired t-map was recomputed from the resulting difference images, and the top-ranked voxels were selected and clustered exactly as for the observed data. Because this null recomputes the statistic from the same images, it preserves the spatial autocorrelation, the noise structure, and the selection procedure of the real data, and removes only the association with orientation.

The second, a scattered-voxel null, is more liberal but makes fewer assumptions. Here the same number of selected voxels was placed at random within each stage mask and clustered, removing spatial structure altogether.

#### Matched-patch sampling

To control for differences in extent between processing stages, we additionally compared each target stage against patches drawn from earlier stages and matched on two quantities: patch volume and the number of selected voxels within the patch. Patches were grown from a randomly chosen seed voxel within the source stage mask until they reached the volume of the target stage; patches not reaching the exact target volume were rejected and redrawn. Within each accepted patch, the same number of voxels as selected in the target stage was chosen by highest absolute t-statistic, and these were clustered using the procedure above. We drew 10,000 patches per source stage. We compared the late stage against patches from the early and mid stages, and the mid stage against patches from the early stage (Fig. 2E). In the latter comparison, mid-stage voxels were selected at p < 0.05 within the mid stage, rather than at the late-stage proportion used above. Statistical significance was assessed by two-sided permutation test, computed as twice the smaller of the two one-tailed proportions of null samples at or beyond the observed value, with the observed value included in the count; the smallest attainable p is therefore 2 × 10⁻⁴.

#### Orientation organization in multivoxel representation space

To quantify how orientation information is distributed within the multivoxel representational space, we devised and implemented a stepwise incremental decoding procedure across visual areas from V1 to IT. We first sorted voxels in descending order based on the absolute value of their orientation contrast t-statistic. Starting with the 10 most informative voxels, we incrementally added voxels one at a time, recalculating decoding accuracy at each step as the dimensionality, and hence participating proportion of each area, increased.

This resulted in a decoding accuracy curve as a function of the proportion of voxels participating in classification per brain area. We defined the multivariate metric, peakProp, as the proportion of voxels at which this curve reached its maximum. The curve was first smoothed with a Gaussian kernel (window = 10 steps) and rescaled between 0.5 and 1, after which peakProp was computed. We computed peakProp using a centroid definition, in which the peak is the weighted center of mass of the upper region of the curve, with weights max(0, c − q) where q is the 90th percentile of the smoothed, rescaled curve.

In parallel, we computed the univariate metric, sigProp, as the proportion of voxels in each ROI reaching p < 0.05 in the univariate orientation contrast. We defined the information distribution distance as:

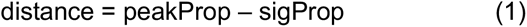

Larger distances indicate more distributed representations, in which collectively informative patterns arise from voxels that are not individually informative.

Confidence intervals on peakProp and on the distance were obtained by bootstrapping. The decoding step returns one accuracy value per classifier repetition at each dimensionality; rather than averaging these away, we retained all 100 repetitions and resampled them with replacement 1,000 times, recomputing the mean curve and re-estimating the peak on each resample. The reported interval is the 2.5th to 97.5th percentile of the resulting distribution. This requires no additional classifier training and reflects variability in the peak estimate rather than in decoding accuracy per se.

When comparing the distance in IT against matched samples from V1, the distance metric could in principle be inflated in IT for two reasons unrelated to representational format. First, V1 is more than twice the size of IT in both animals, so any given voxel proportion in V1 corresponds to a far larger number of voxels, and the V1 decoding curve may reach its maximum at a lower proportion simply because that proportion already affords a high-dimensional pattern. Second, given the placement of the receive coils, contrast t-statistics are likely inflated in V1 relative to anterior temporal cortex, so that fewer IT voxels pass threshold and sigProp is correspondingly lower.

To address both, we constructed two null distributions from subsamples of V1, each matched to IT and each of 200 samples. In the first (V1size), voxels were drawn from V1 at random to match IT’s voxel count. In the second (V1sel), voxels were additionally matched to IT’s distribution of absolute t-statistics: we binned IT’s |t| values into 20 quantile bins and drew, from each bin, the same number of V1 voxels as IT contained, filling any deficit from the remaining unselected pool. Because this matching reproduces IT’s univariate selectivity profile, sigProp is equated between the sample and IT by construction, and any residual difference in the distance cannot be attributed to differences in univariate selectivity or signal quality.

For each subsample, the incremental decoding procedure was rerun within the sampled voxel set alone, with voxel ranking restricted to that set, and peakProp, sigProp and their distance recomputed exactly as for the observed data (400 dimensionality steps, 10 classifier repetitions per step). The observed IT distance was compared against each null as

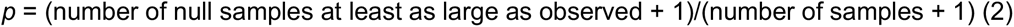

#### Connectivity between orientation-representing voxels

Connectivity was computed on the blank intervals between stimulus blocks to minimize stimulus-driven effects such as co-activation. For each run, we first removed low-frequency drift by regressing out a second-order polynomial (constant, linear and quadratic terms) from each voxel’s full timeseries. We then identified the blank blocks from the paradigm file and discarded the first 5 TRs (10 s) following stimulus offset to allow the hemodynamic response to return towards baseline, retaining the remaining 7 TRs (14 s) of each block. Each retained block was demeaned and linearly detrended independently, so that residual differences in signal level or slope between blocks could not contribute to the correlation across concatenated blocks. Finally, each voxel’s concatenated blank timeseries was divided by its own standard deviation, equating the scale across voxels. Blocks were then concatenated within and across runs, yielding a single blank-period timeseries per voxel on which all connectivity measures were computed.

To measure functional connectivity, we computed Pearson correlation coefficients between all voxel pairs on the blank-period timeseries described above, and converted them to Fisher z (correlations were clipped to ±(1 − 10⁻⁷) before transformation to avoid infinite values). We grouped the resulting pairwise connectivity values into four categories:

i. **top2top**: between significant voxels in both upstream and downstream areas
ii. **downTop**: between significant downstream voxels and all upstream voxels
iii. **upTop**: between significant upstream voxels and all downstream voxels
iv. **all**: between all voxels across the two regions

We performed this procedure across three hierarchical processing stage pairs: early-mid, early-late, and mid-late. We then plotted the average connectivity across the four categories (top2top → downTop → upTop → all) to evaluate whether orientation responsiveness systematically influenced inter-areal functional connectivity.

Voxel pairs are not independent: each voxel contributes to as many pairs as there are voxels in the opposing region, so treating pairs as independent observations would substantially overstate the precision of the resulting means. We therefore based all inference on a bootstrap over voxels rather than over voxel pairs. In each of 10,000 iterations, upstream and downstream voxels were independently resampled with replacement, and the mean connectivity for each of the four conditions was recomputed over the pairs formed by the resampled sets. Because voxels are the resampling unit, all pairs sharing a voxel enter or leave a given resample together, and the resulting distribution reflects variability across voxels rather than across pairs. Error bars in Fig. 3B are the standard deviation of this bootstrap distribution.

Each of the three feature-specific conditions was compared against the whole-region condition (*all*) by taking the difference of their bootstrap distributions and computing a two-sided p as twice the smaller tail proportion, with a floor of 1/10,000. As a summary of the overall trend, we additionally report the proportion of bootstrap resamples in which the complete ordering top2top > downtop > uptop > all was preserved. Effect sizes are reported as Cohen’s d.

To test robustness of these connectivity findings, we repeated this analysis across all 15 pairwise combinations of individual visual ROIs (e.g., V1-V2, V1-V3, etc.). Similar to the stage-wise analysis, we compared connectivity among significant voxels to a null distribution obtained through size-matched random sampling from non-significant voxels over 1000 permutations using unpaired t-tests.

Finally, to determine the specificity of the observed connectivity effects to visual areas, we repeated this procedure for all possible pairwise combinations of visual and control ROIs (e.g., V1-MT, V1-LIP, etc.) and compared the results with exclusively visual pairs obtained above (see Supplementary Fig. 1).

#### Information transformation along the visual hierarchy

To investigate information transfer across the visual hierarchy, we combined connectivity and decoding approaches. For each significant upstream voxel (based on contrast t-statistic, p < 0.05), we identified its most strongly connected downstream voxel. This process was repeated for all significant upstream voxels, with the constraint that each downstream voxel could be assigned to only one upstream voxel. If a downstream voxel had already been assigned, we selected the next strongest unassigned voxel, ensuring one-to-one mapping between upstream and downstream voxels.

We named the two corresponding voxels top, and top-connected in upstream and downstream respectively. We performed this assignment across all 15 possible pairs of visual regions along the hierarchy (i.e., V1–V2, V1–V3, V1–V4, V1–TEO, V1–TE, V2–V3, …, TEO–TE).

To quantify how much information was shared between top and top-connected voxels, we trained a linear SVM classifier on the top upstream voxels and tested it on the corresponding top-connected downstream voxels. The higher cross-region decoding accuracy indicated greater similarity in the geometry of the representation space, reflecting stronger information transfer.

To test statistical significance of the observed transfer decoding performance, we again turned to random sampling and permutation approach: we randomly selected voxel subsets from upstream and downstream areas (matched in size to the top voxels set), trained an SVM on the upstream sample, and tested it on downstream sample. This random sampling and classification was repeated for 1000 permutations to generate a null distribution. We then statistically tested the observed transfer accuracy against this null distribution using unpaired t-tests for each region pair. This analysis was repeated independently for all 15 visual area pairs (see Fig. 3D).

#### Clustering in Face-Object-Body (FOB) space and corresponding orientation information

As for the analyses of orientation information, we first regressed out nuisance variables from the Face-Object-Body (FOB) localizer data. We then averaged BOLD activity during face, body, and object presentations, respectively. We subtracted the same-run blank background baseline to compute evoked responses. We then z-scored each voxel’s evoked response separately for each category condition, using the mean and standard deviation computed across all voxels within the respective region of interest (e.g., TE, etc.). Voxels with a z-score exceeding a threshold of 1.2 were marked as responsive to the respective category (results were robust across a range of reasonable thresholds; see Supplementary Fig. 7A for details). Voxels were categorized based on their response profiles into one of the following groups:

- **Mix**: responsive to all three categories
- **Double**: responsive to any two of the three categories
- **Pure**: responsive to only one category
- **Non-responsive**: bottom 20% of responses across all conditions

We performed this classification independently within areas V4, TEO, and TE. V4 was included as a mid-level control region, while TEO and TE were selected for their established roles in high-level object category representation (Kravitz et al., 2013).

We then computed orientation information for each voxel in these regions using the absolute contrast t-statistic derived from the orientation mapping data. Orientation information distributions were compared across the defined FOB clusters (non-responsive, pure, double, and mix). Results from TEO and TE were combined to represent area IT and are shown in Fig. 4C. Individual area results are provided in Supplementary Fig. 7B.

#### Mix clusters vs. the rest of IT

To test whether mix clusters are more strongly connected to upstream areas than the remaining IT population, we compared them with size-matched random sets of non-mix IT voxels. The non-mix pool comprised all IT voxels outside mix clusters. We drew 5,000 random sets from this pool, each containing the same number of voxels as the mix clusters and drawn without replacement. The same 5,000 sets were used for all upstream areas and for both analyses below, so that the two analyses are directly comparable.

Connectivity to upstream areas (Fig. 5C): For each IT voxel i, we computed its connectivity to a given upstream area, c_i, as the mean Fisher z-transformed correlation between voxel i and all voxels of that area. The observed value was the mean of c_i over mix voxels; each random set yielded one value of the null distribution, the mean of c_i over the voxels of that set.

Preference for orientation-selective upstream voxels (Fig. 5D): Within each upstream area, voxels were divided into orientation-informative (p < 0.05 on the orientation contrast) and non-informative (p > 0.05) groups. For each IT voxel i, the preference was defined as Δc_i = c_i(informative) − c_i(non-informative), where c_i(informative) is the mean connectivity of voxel i to all informative voxels, and c_i(non-informative) is its mean connectivity to a random sample of non-informative voxels matched in number to the informative group, averaged over 1,000 such samples. To summarize the preference of a group of IT voxels, we used:

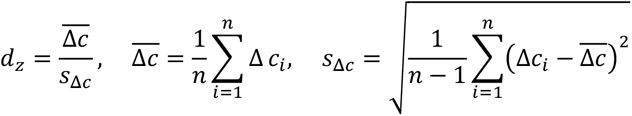

where Δ*c_i_* the preference of IT voxel i, i.e., its connectivity to orientation-informative upstream voxels minus its connectivity to non-informative upstream voxels; *Δc* is the mean of Δ*c_i_*over the n voxels of the group (mix clusters or a random set of non-mix voxels); *s*_Δ*c*_ is the corresponding standard deviation. *d_z_* thus expresses how consistently the voxels of a group are more strongly connected to informative than to non-informative upstream voxels.

We computed d_z for the mix voxels and for each of the 5,000 random sets, and formed the distribution of d_z(mix) − d_z(random set).

In both analyses, p values are one-sided (mix > rest of IT), computed as the proportion of random sets with a value at least as large as that of the mix clusters, with a floor of 1/5,000 (for Fig. 5D, this equals the proportion of the d_z difference distribution at or below zero). P values were FDR-corrected (Benjamini–Hochberg) across the four upstream areas. Effect sizes are reported as Cohen’s d,

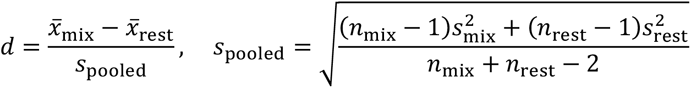

where *x̄*_mix_ and *x̄*_rest_ are the means of the per-voxel values (*c_i_* in Fig. 5C, Δ*c_i_* in Fig. 5D) over mix voxels and over all remaining IT voxels, 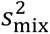 and 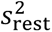 are the corresponding variances, and *n*_mix_ and *n*_rest_ are the numbers of voxels in each group.

Cohen d compares the per-voxel values of mix voxels with those of all remaining IT voxels. 95% confidence intervals for d were obtained from 5,000 bootstrap resamples of voxels, drawn with replacement within each group (2.5th and 97.5th percentiles). Confidence intervals of the mean mix connectivity were obtained by a subsampled pair bootstrap: in each of 5,000 replicates, 90% of the contributing voxel pairs were drawn without replacement and their mean recorded, and the standard deviation of these means was rescaled by 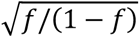 where f is the subsampling fraction, to recover the full-sample standard error (intervals: mean ± 1.96 rescaled SD).

#### Connectivity pure vs. mix clusters

As a more specific control, pure and mix clusters were compared directly using the same connectivity measure (c_i), preference measure (Δc_i, d_z) and effect size (d) as in Fig. 5C, D. For connectivity to upstream areas (Supplementary Fig. 5A), each cluster type was compared with size-matched random sets of the IT voxels outside pure and mix clusters (5,000 sets), and the Mix − Pure difference was tested against 10,000 shuffles of the pure and mix labels. For connectivity to orientation-selective upstream voxels (Supplementary Fig. 5B), the mean connectivity of each cluster type to informative voxels was compared with a null of 1,000 size-matched samples of non-informative voxels, and the Mix − Pure difference was tested against the difference of the two nulls. Per-class p values are two-sided and Mix − Pure p values one-sided (mix > pure); all were FDR-corrected across the four upstream areas. For the Mix − Pure contrast, d compares the per-voxel values of mix and pure voxels.

#### DCNN Models’ response to Face–Object–Body (FOB) categories and orientation

To test whether the enhanced orientation information observed in mix voxels is replicated in artificial neural networks, we analyzed a class of deep convolutional neural networks (DCNNs) selected based on their similarity to the primate visual system using the Brain-Score metric (Schrimpf et al., 2020). This metric quantifies the similarity of DCNNs to the primate brain across multiple dimensions—including perceptual behavior and representational similarity— and provides an overall score indicating model–brain correspondence. We selected 17 DCNN architectures with a Brain-Score above 0.5 (results were robust to more stringent thresholds; see Fig. 6B for a complete list of models). This process was intended to ensure the inclusion of the most relevant models for comparison with the orientation–cluster effect observed in the monkey’s brain.

The DCNNs included in this study were pretrained on the ImageNet dataset, which contains over 1.2 million images across 1000 object categories, and were implemented using the PyTorch framework.

To examine whether these models replicate the orientation–category interaction observed in the brain, we applied the same analysis used in fMRI voxel space to the models’ penultimate layer, chosen for its correspondence to IT cortex as the final stage of visual representation in the brain.

We defined pure, double, and mix units in each model using the same clustering logic as applied to IT voxels — based on their category-specific evoked responses to faces, objects, and bodies. Orientation information for each unit was quantified as the absolute difference in response to the two diagonal grating stimuli. This procedure was performed separately for each model.

To test for any systematic differences in orientation information across category clusters, we first averaged each model’s orientation information within each cluster. We then compared these model-averaged values across cluster pairs (mix vs. pure, mix vs. double, double vs. pure) using two-sided Wilcoxon signed-rank tests, paired across the 17 models, and FDR-corrected (Benjamini–Hochberg) across the three comparisons.

Finally, to quantify connectivity between either of clusters in penultimate layer, analogous to fMRI connectivity analysis, we turned to upstream influence analysis. upstream influence was defined using a gradient-based sensitivity measure, following approaches in deep network interpretability (Bach et al., 2015; Simonyan et al., 2014; Sundararajan et al., 2017) and receptive-field analysis (Alain & Bengio, 2018; Luo et al., 2016). This measure computes the total leverage of upstream activations on each penultimate unit, consistent with gradient-based definitions of unit importance widely used in machine learning and neuroscience (Cadieu et al., 2014; Yamins & DiCarlo, 2016). Specifically, for each penultimate unit *v_d_*, we define its upstream influence as:

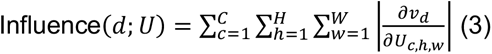

Where *U_c_*_,*h*,w_ is the activation tensor of the upstream layer (channels × height × width), *v_d_* is the scalar activation of the *d-th* penultimate unit.

This quantity captures the local, linearized sensitivity of penultimate unit *v_d_* to infinitesimal perturbations anywhere in the upstream feature map *U_c_*_,*h*,w_. Taking the absolute value and summing across all channels and spatial locations yields a global, sign-agnostic measure of total upstream influence on that unit.

To apply the same mix vs. size-matched non-mix comparison as in the fMRI data, we compared, for each architecture and each candidate V1-, V2-, and V4-like layer (see Supplementary Table 2), the mean upstream influence of mix units with that of size-matched random sets of non-mix units (10,000 sets), following the sampling procedure described for Fig. 5C. The non-mix pool comprised non-responsive, pure, and double-responsive units; units with zero influence (inactive for the probe input) were excluded. To account for differences in scale across architectures, the observed value was expressed relative to its null as

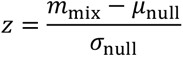

where *m*_mix_ is the mean influence of mix units, and *μ*_null_ and *σ*_null_ are the mean and standard deviation of that model’s null. Across the 17 models, the median z was compared with a null of the median, obtained by taking, for each random set, the median of the models’ standardized null values. P values are two-sided and FDR-corrected across the three layers.

In addition, as for the fMRI data, we compared cluster types directly: the average upstream influence of pure, double, and mix units was z-scored within each architecture across cluster types and compared across the 17 models with the Wilcoxon signed-rank test (Supplementary Fig. 5C).

#### Extracting and comparing orientation content of object categories

To control for potential confounds arising from systematic differences in orientation content across stimulus categories (faces, bodies, and objects), we quantified and compared the orientation content of all visual stimuli using Gabor filters (Adelson & Bergen, 1985; Jones & Palmer, 1987; Kay et al., 2008; Movellan, 2002). The goal was to assess whether low-level image features could account for the observed relationship between IT orientation sensitivity and object selectivity.

We defined the 2D Gabor filter as:

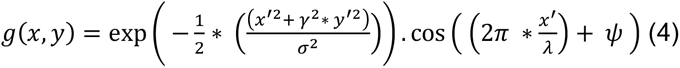

With rotated coordinates (*x*^′^ , *y*′):

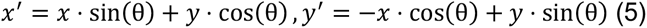

Where:

*σ* is standard deviation of Gaussian envelope,

θ is orientation of the Gabor filter in radians,

*λ* is wavelength of the sinusoidal factor in pixels,

*γ* is spatial aspect ratio, and

*ψ* gives phase offset of the sinusoidal wave.

For each image, we constructed two Gabor filter banks: one to capture fine (local) orientation content and another to capture coarse (global) orientation content. This was achieved by modifying the filter parameters. For fine orientation detection, we used *size*(*x*, *y*) = 100, *γ* = 0.9 and *λ* = 5; for coarse orientation, we used the full image size, *γ* = 0.5 and *λ* = 10 for detecting coarse (global) orientation. These values were chosen based on visual inspection of filter responses, and results were robust across a range of reasonable parameter settings.

This way, we computed average fine and coarse orientation contents for each images. We then compared the distributions of orientation content across stimulus categories and against the two diagonal grating conditions using circular statistical tests implemented in the Circular Statistics Toolbox (Berens, 2009). Specifically, we applied the Watson–Williams multi-sample test to assess differences in mean orientation content across object categories, and the V-test to evaluate whether object orientations were non-uniformly distributed and biased toward the two diagonal orientations.

## Data and code availability

The data and analysis code that support the findings of this study are available upon reasonable request from the corresponding author.

