## Supplemental Information for "Re-emergence of orientation coding in primate IT cortex and deep networks reveals functional hubs for visual processing"

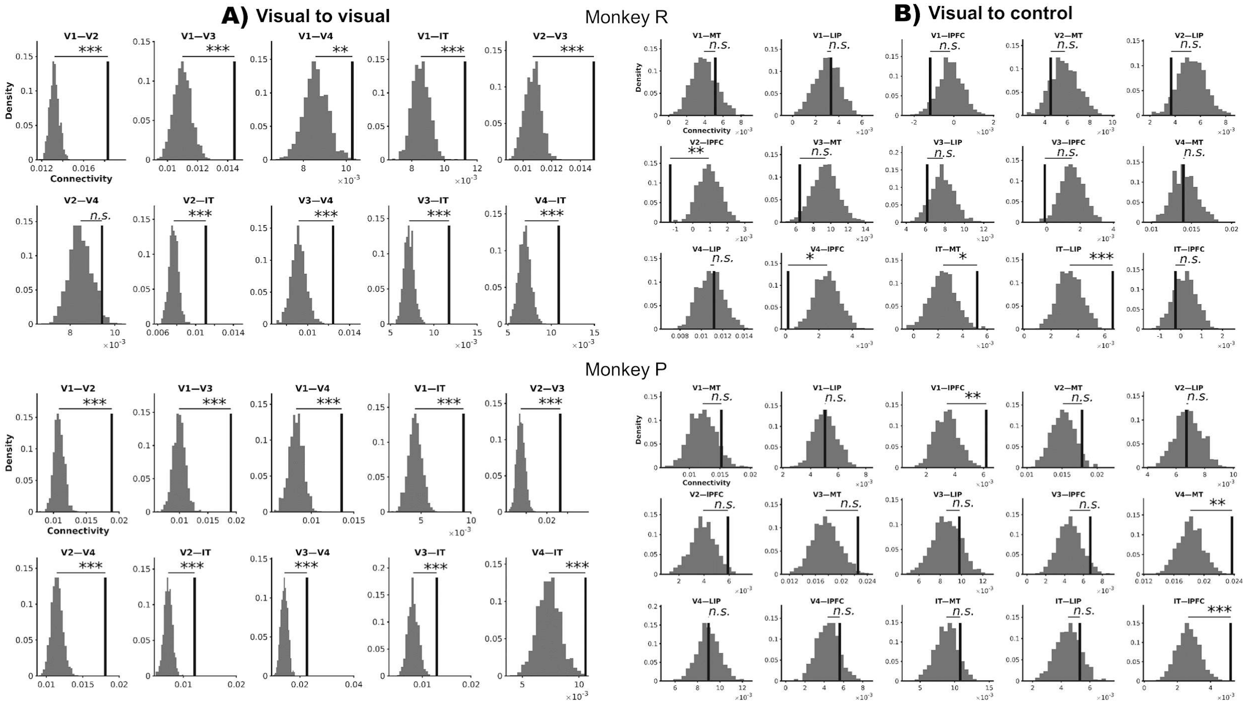


**Supplementary Figure 1.** *Connectivity between pairs of visual areas and between visual and control areas in two monkeys.* **A.** Histograms of connectivity estimates (z-transformed Pearson correlation) for all pairwise combinations of visual areas (V1, V2, V3, V4, IT) in monkeys R and P, respectively. Vertical black lines indicate observed connectivity, and gray distributions show null distributions obtained from size-matched permutations. Statistically significant connectivity is observed across nearly all visual–visual pairs (**, $p < 0.01$; ***, $p < 0.001$; n.s., not significant). **B.** Same analysis for connections between visual areas and control regions (MT, LIP, IPFC). Most visual–control pairs do not reach statistical significance, with only a subset showing weak connectivity (**, $p < 0.01$; ***, $p < 0.001$; n.s., not significant).

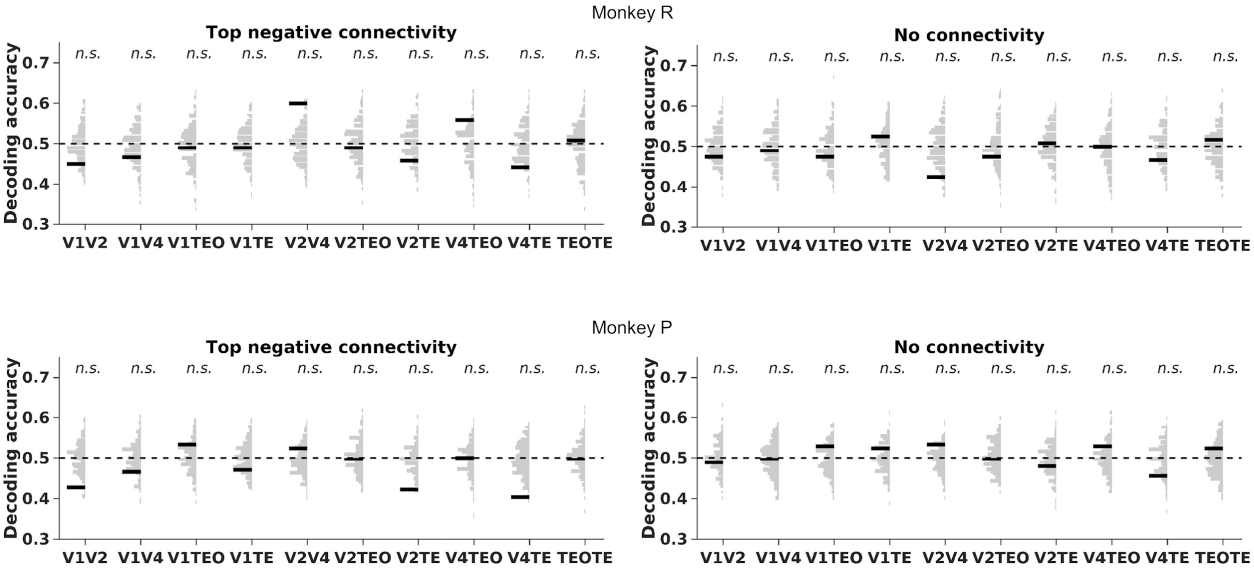


**Supplementary Figure 2.** *Control analyses of representational transfer using negatively connected and unconnected voxel pairs.* To assess representational transfer under alternative connectivity definitions, top voxels in upstream regions were mapped one-to-one to their weakest connected voxels in downstream regions (top negative connectivity; smallest Pearson correlations) or to voxels with near-zero connectivity (no connectivity). A linear SVM classifier was trained on multivoxel patterns from upstream voxels and tested on the corresponding downstream voxels in each control group. Decoding accuracy is shown for all upstream–downstream pairs in monkeys R (top) and P (bottom). Thick black bars indicate observed transfer accuracies, and light gray bars represent null distributions generated by random size-matched voxel sampling for statistical comparison. The dashed black line marks chance level (0.5). No upstream–downstream pair reached statistical significance ($p < 0.05$) under these control conditions.

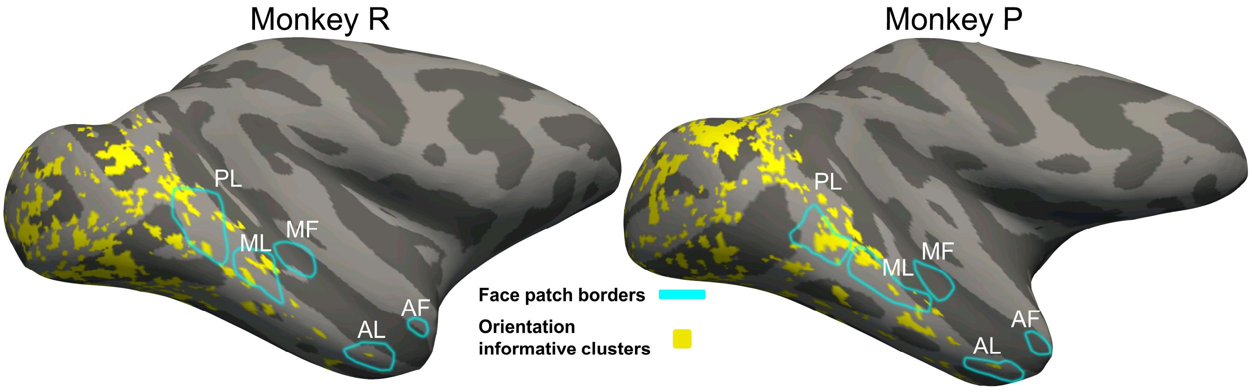


**Supplementary Figure 3.** *Spatial relationship between BOLD-defined face patches and orientation-informative voxels in the right hemisphere.* Inflated right-hemisphere surfaces are shown for Monkey R and Monkey P. Cyan outlines indicate the borders of face-selective patches identified from the localizer contrast of faces versus all other object categories, including man-made objects, monkey bodies, vegetables and fruits, and phase-scrambled images. Posterior face patches, including PL, ML, and MF, were delineated using a threshold of $p < 10^{-5}$, corresponding to $t > 4.4$, whereas anterior face patches, including AL and AF, were delineated using a threshold of $p < 0.05$, corresponding to $t > 1.96$. Yellow overlays indicate orientation-informative voxels projected onto the same inflated surfaces. The presence of clearly identifiable face-patch organization demonstrates that face-selective patches can be identified from BOLD signal alone using the procedures described in the Methods, without the use of a MION contrast agent. Across both animals, orientation-informative voxels do not show consistent colocalization with face-patch borders or a systematic relationship with the anterior–posterior face-patch organization.

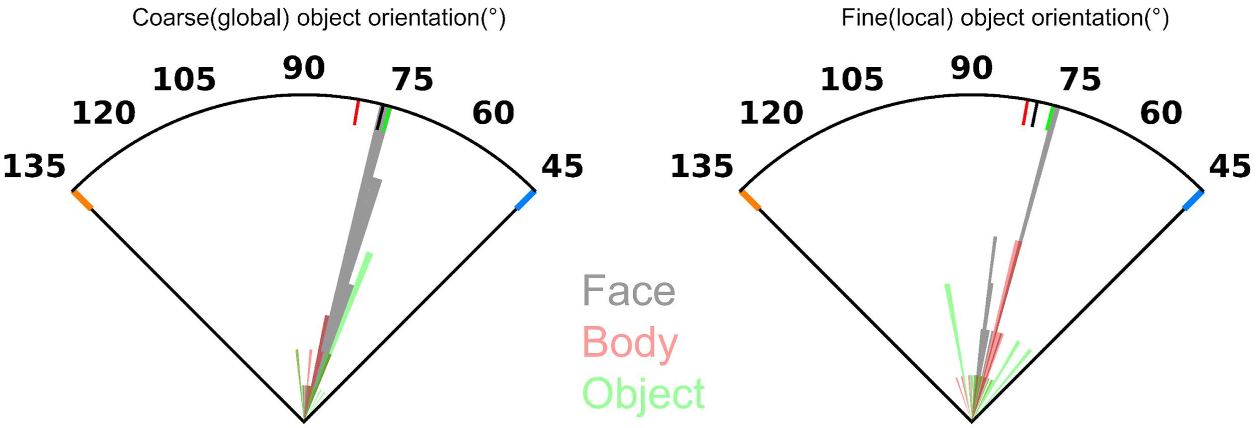


**Supplementary Figure 4.** *Distribution of coarse and fine object orientations across categories.* Polar plots illustrate the orientation structure of images from three object categories (faces, bodies, and objects). Orientations were computed at two levels (see Methods): coarse (global) object orientation, defined as the dominant axis of the whole object silhouette (left), and fine (local) object orientation, defined from local edge filters applied within object boundaries (right). For each category, light-colored lines show the polar distribution of orientations across individual exemplars, while dark-colored lines indicate the average observed orientation for that object. Colors correspond to object categories (gray = face, red = body, green = object). Orientation distributions were significantly biased relative to both 45° and 135°, but did not differ significantly from each other ($p < 0.05$). Tick marks denote orientation angles in degrees.

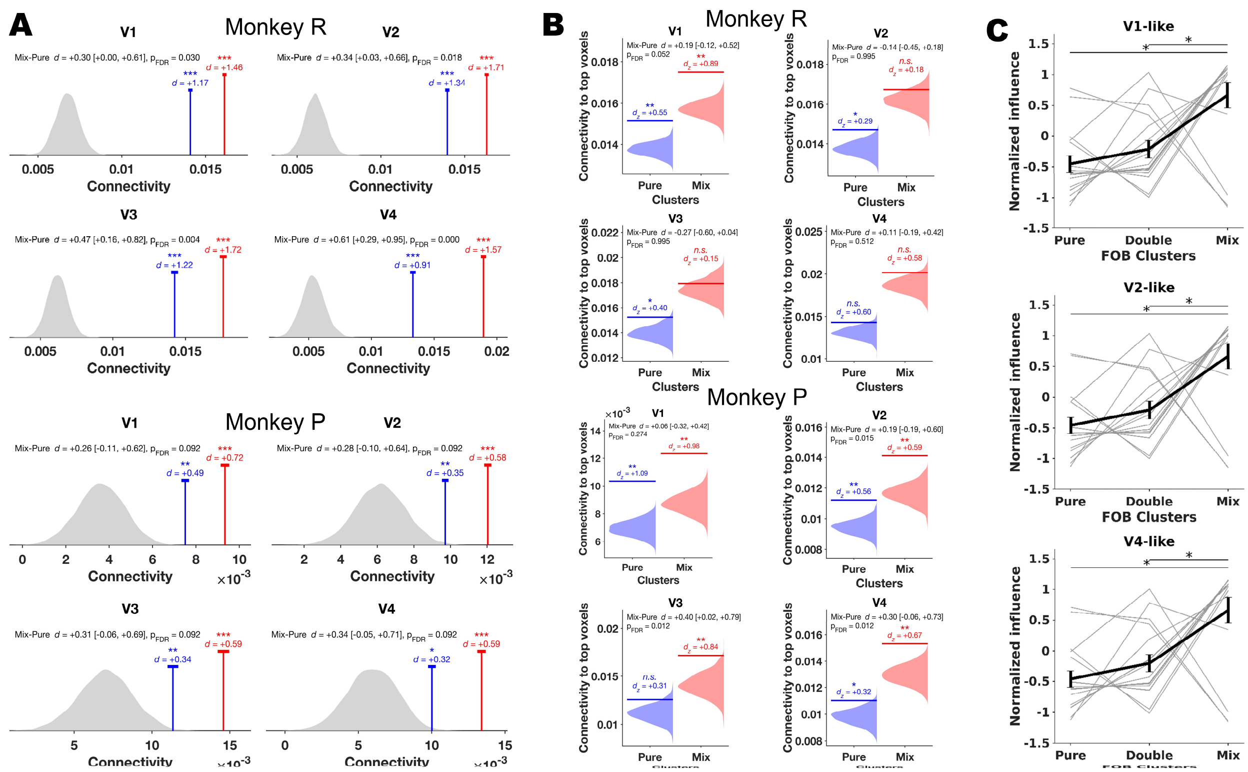


**Supplementary Figure 5.** *Retinotopic mapping of polar angle and eccentricity in visual cortex.* Polar angle (top) and eccentricity (bottom) maps are shown on inflated lateral surfaces of monkeys R and P, respectively. Maps are restricted to visual cortex, and dashed lines mark borders of visual areas as defined by polar angle reversals. The color wheels indicate the polar angle and eccentricity conventions used for mapping.

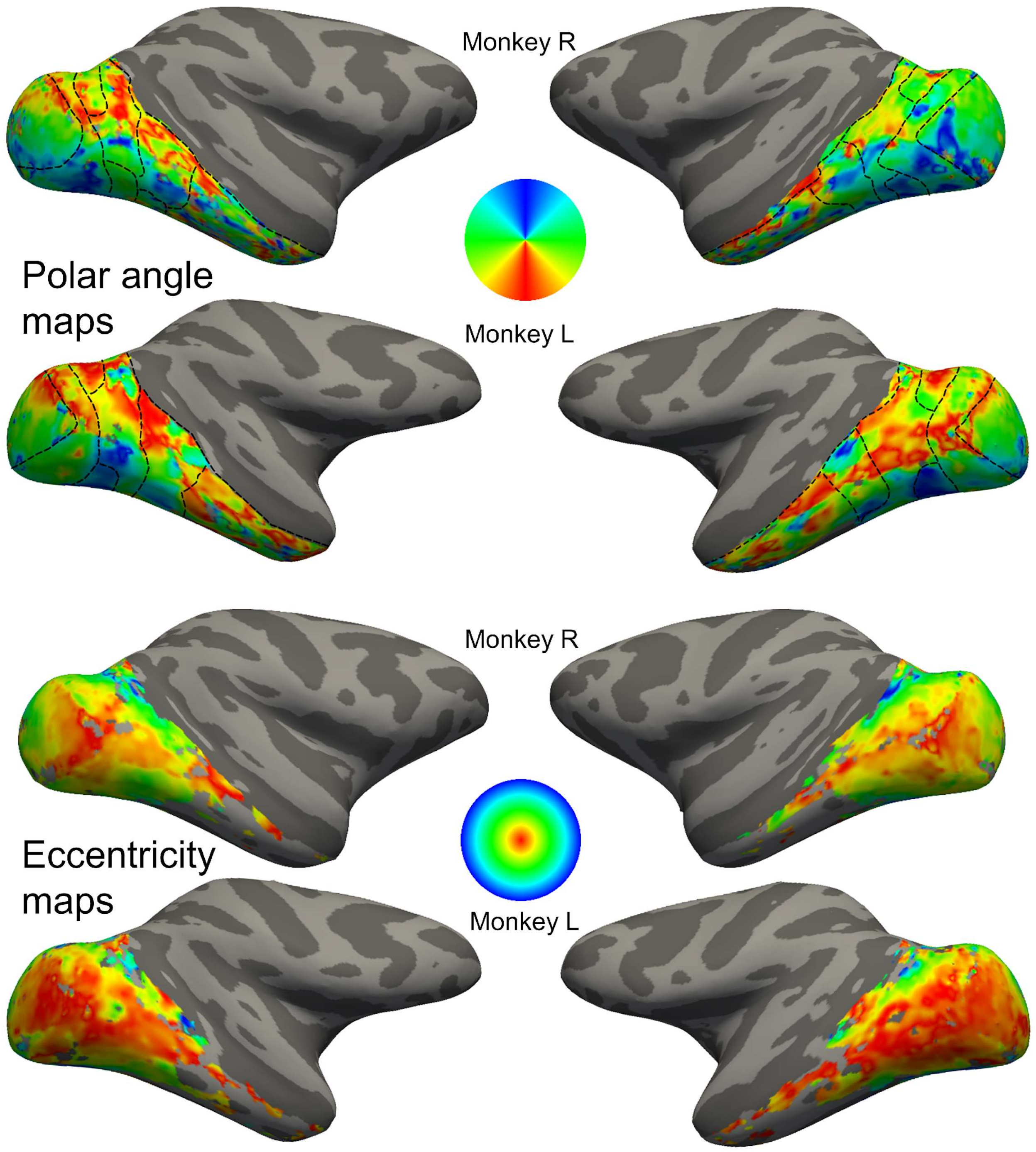


**Supplementary Figure 6.** *Retinotopic mapping of polar angle and eccentricity in visual cortex.* Polar angle (top) and eccentricity (bottom) maps are shown on inflated lateral surfaces of monkeys R and P, respectively. Maps are restricted to visual cortex, and dashed lines mark borders of visual areas as defined by polar angle reversals. The color wheels indicate the polar angle and eccentricity conventions used for mapping.

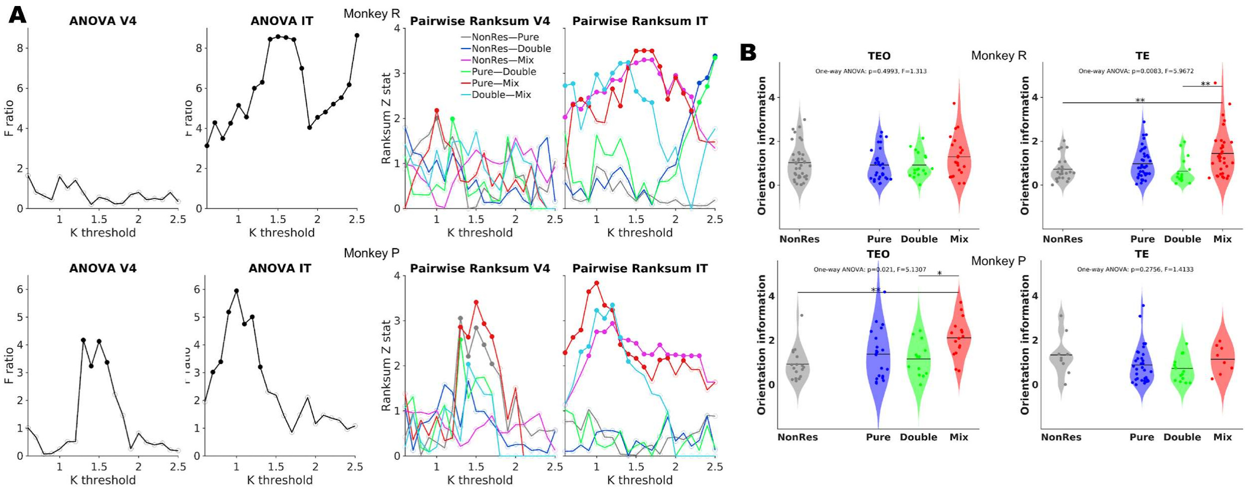


**Supplementary Figure 7.** *Robustness of responsive voxel thresholding and cluster-type selection.* **A.** One-way ANOVA *F*-ratios (left) testing the effect of voxel cluster type (NonRes, Pure, Double, Mix) on orientation information, and pairwise rank-sum statistics (right) across a range of voxel responsiveness thresholds (*K*) in V4 and IT for monkeys R (top) and P (bottom). Here, *K* indicates the z-threshold used to classify voxels as responsive, which was later used to define voxel types (Pure = responsive to only one category, Double = responsive to two of the three categories, Mix = responsive to all three categories: face, body, and object). Filled circles denote threshold levels at which ANOVA *F*-ratios or rank-sum Z-statistics reached statistical significance, while open circles indicate nonsignificant values. Results confirm that findings are robust across thresholds, with IT showing stronger and more consistent separation among voxel types compared to V4. **B.** Violin plots showing the distribution of orientation information (quantified as the absolute value of the orientation contrast *t*-statistic) across the four voxel types defined in A, shown separately for TEO and TE in each monkey. Black horizontal lines indicate medians, and shaded regions denote standard deviations. One-way ANOVA results are indicated on each panel; pairwise comparisons were conducted, and only significant results are shown. ***, $p < 0.001$; **, $p < 0.01$; *, $p < 0.05$; n.s., not significant. These individual-area results complement Fig. 4C, where TEO and TE were combined into IT.

**Table 1 – GLM results for predicting orientation response using object category response**

| **Monkey R – V4** | | | | |
|---|---|---|---|---|
| Term | Beta | SE | tStat | pValue |
| face | 0.0025697 | 0.019916 | 0.12902 | 0.89747 |
| body | 0.069829 | 0.044282 | 1.5769 | 0.11639 |
| obj | 0.0064226 | 0.016121 | 0.3984 | 0.69076 |
| face×body | -0.00044862 | 0.0073562 | -0.060986 | 0.95143 |
| face×obj | -0.00036269 | 0.0024291 | -0.14931 | 0.88146 |
| body×obj | -0.0051974 | 0.0075534 | -0.68809 | 0.49219 |
| face×body×obj | 0.00021665 | 0.00081327 | 0.26639 | 0.79021 |
| **Monkey R – IT** | | | | |
| Term | Beta | SE | tStat | pValue |
| face | -0.0042522 | 0.0099406 | -0.42776 | 0.66931 |
| body | 0.020851 | 0.031069 | 0.67113 | 0.50295 |
| obj | 0.013823 | 0.020306 | 0.68072 | 0.49687 |
| face×body | 0.0020205 | 0.0029471 | 0.68558 | 0.4938 |
| face×obj | -0.00041453 | 0.0025603 | -0.16191 | 0.87155 |
| body×obj | -0.004582 | 0.0051933 | -0.88229 | 0.37872 |
| face×body×obj | 0.00025553 | 0.00057355 | 0.44552 | 0.65645 |
| **Monkey P – V4** | | | | |
| Term | Beta | SE | tStat | pValue |
| face | 0.025318 | 0.03024 | 0.83722 | 0.40386 |
| body | 0.079962 | 0.053275 | 1.5009 | 0.13557 |
| obj | 0.0275 | 0.043846 | 0.62719 | 0.53153 |
| face×body | -0.011816 | 0.014337 | -0.82417 | 0.41121 |
| face×obj | -0.0046412 | 0.015325 | -0.30285 | 0.76244 |
| body×obj | -0.0050684 | 0.022291 | -0.22737 | 0.82046 |
| face×body×obj | -0.0032685 | 0.0061663 | -0.53006 | 0.59688 |
| **Monkey P – IT** | | | | |
| Term | Beta | SE | tStat | pValue |
| face | 0.032087 | 0.026283 | 1.2209 | 0.22369 |
| body | -0.062495 | 0.047127 | -1.3261 | 0.18644 |
| obj | -0.047564 | 0.040576 | -1.1722 | 0.24262 |
| face×body | 0.0057677 | 0.0070004 | 0.82391 | 0.41105 |
| face×obj | 0.012879 | 0.0071661 | 1.7972 | 0.073935 |
| body×obj | 0.025185 | 0.013819 | 1.8225 | 0.069987 |
| face×body×obj | -0.00030227 | 0.00082362 | -0.367 | 0.71404 |

**Table 2 – layer properties for each individual DCNN**

| Model(s) | V1-like layer | V2-like layer | V4-like layer | Penultimate layer |
|---|---|---|---|---|
| DN169 (DenseNet-169) | features[0] (conv0) | features[4] (denseblock1) | features[6] (denseblock2) | features[10] (denseblock4) |
| CorNet (CORnet-S) | mdl.V1 | mdl.V2 | mdl.V4 | mdl.IT (pre-classifier) |
| RN101 (ResNet-101) | layer1 | layer2 | layer3 | layer4 (last conv stage) |
| DN201 (DenseNet-201) | features[0] (conv0) | features[4] (denseblock1) | features[6] (denseblock2) | features[10] (denseblock4) |
| DN121 (DenseNet-121) | features[0] (conv0) | features[4] (denseblock1) | features[6] (denseblock2) | features[10] (denseblock4) |
| RN152 (ResNet-152) | layer1 | layer2 | layer3 | layer4 (last conv stage) |
| RN50 (ResNet-50) | layer1 | layer2 | layer3 | layer4 (last conv stage) |
| timm (inception_v4, inception_resnet_v2, xception, nasnetalarge) | feature_info[0].module | feature_info[1].module | feature_info[2].module | feature_info[-1].module |
| mobilenetv3_large | features[2] (fallback: features[0] if <3) | features[6] (fallback: ~1/3 of features) | features[12] (fallback: ~2/3 of features) | features[-1] (final block) |
| resnet18 / resnet34 | layer1 | layer2 | layer3 | layer4 (last conv stage) |
| vgg16 / vgg19 | features[0] (conv1_1) | features[5] (conv2_1) | features[10] (conv3_1) | features[28] (conv5_3, pre-ReLU) |
| inception_v3 | Mixed_5d | Mixed_6e | Mixed_7b | Mixed_7c |
